# KAHRP dynamically relocalizes to remodeled actin junctions and associates with knob spirals in *P. falciparum*-infected erythrocytes

**DOI:** 10.1101/2021.05.12.443751

**Authors:** Cecilia P. Sanchez, Pintu Patra, Shih-Ying Chang, Christos Karathanasis, Lukas Hanebutte, Nicole Kilian, Marek Cyrklaff, Mike Heilemann, Ulrich S. Schwarz, Mikhail Kudryashev, Michael Lanzer

## Abstract

The knob-associated histidine-rich protein (KAHRP) plays a pivotal role in the pathophysiology of *Plasmodium falciparum* malaria by forming membrane protrusions in infected erythrocytes, which anchor parasite-encoded adhesins to the membrane skeleton. The resulting sequestration of parasitized erythrocytes in the microvasculature leads to severe disease. Despite KAHRP being an important virulence factor, its physical location within the membrane skeleton is still debated, as is its function in knob formation. Here, we show by super-resolution microscopy that KAHRP initially associates with various skeletal components, including ankyrin bridges, but eventually co-localizes with remnant actin junctions. We further present a 35Å map of the spiral scaffold underlying knobs and show that a KAHRP-targeting nanoprobe binds close to the spiral scaffold. Single-molecule localization microscopy detected ∼60 KAHRP molecules per knob. We propose a dynamic model of KAHRP organization and a function of KAHRP in attaching other factors to the spiral scaffold.

## Introduction

Red blood cells infected with the human malaria parasite *Plasmodium falciparum* acquire thousands of small protrusion that render their initially smooth surface bumpy (Gruenberg et al., 1983). These protrusions, termed knobs, play a pivotal role in the pathophysiology of falciparum malaria. They form a platform on which parasite-encoded adhesins, such as the immune-variant PfEMP1 antigens, are presented and anchored to the membrane skeleton (Warncke and Beck, 2019). As a result, parasitized erythrocytes attain cytoadhesive properties and sequester in the deep vascular bed of inner organs, which, in turn, can lead to severe sequelae including impaired tissue perfusion, hypoxia and local microvascular inflammation followed by barrier dysfunction (Lee et al., 2019; Smith et al., 2013). Knobs also play an important role in reorganizing and stiffening the cell envelope, leading to rounder shapes and reduced deformability (Fröhlich et al., 2019; Zhang et al., 2015).

Previous studies have suggested that knobs are supramolecular structures composed of parasite-encoded factors and components of the erythrocyte membrane skeleton (Warncke and Beck, 2019). A central building block is the parasite-encoded knob-associated histidine-rich protein (KAHRP). Parasite mutants lacking the corresponding gene are knobless and do not cytoadhere in flow (Crabb et al., 1997). KAHRP features several low affinity interaction domains, allowing it to self-aggregate and to bind to actin, ankyrin, spectrin, and the cytoplasmic domain of PfEMP1 antigens (Cutts et al., 2017; Kilejian et al., 1991a; Oh et al., 2000; Pei et al., 2005; Waller et al., 1999; Warncke and Beck, 2019; Weng et al., 2014). However, spatial and temporal aspects of the assembly are still debated. In particular, there are dissenting views as to where knobs form - at the actin junctional complex (Oh et al., 1997; Oh et al., 2000), the ankyrin bridge (Cutts et al., 2017) or in the mesh formed by spectrin filaments (Looker et al., 2019). In a recent development, it has been shown that knobs are underlaid with a spiral-like scaffold (Watermeyer et al., 2016). However, the molecular composition of this spiral is still unclear, as is the overall architecture of knobs.

Recent advances in super-resolution fluorescence microscopy and image processing (Hell and Wichmann, 1994; Schnitzbauer et al., 2018) have offered new opportunities to interrogate the membrane cytoskeletal organization of erythrocytes at the nanometer scale (Pan et al., 2018). To gain insights into the structural organization of knobs, we implemented several high-resolution imaging techniques, including two-color stimulated emission depletion (STED) microscopy with two-dimensional pairwise cross-correlation analysis, photo-activated localization microscopy (PALM) with single molecule counting, and electron tomography labelling with specific nanoprobes and stereological computer simulations. The combination of these imaging platforms allowed us to map the physical localization of KAHRP within the membrane skeleton and on the spiral scaffold underlying knobs over different stages of parasite development.

## Results

### Analysis of the membrane skeleton of red blood cells by super-resolution microscopy

To investigate the organization of knobs and their interaction with the membrane skeleton, we used a recently developed imaging protocol based on lysing erythrocytes and exposing their plasma membrane on a planar substrate (Fig. 1A) (Looker et al., 2019). The exposed membranes were subsequently stained with antibody combinations, e.g., against the N-terminus (actin-binding domain) of ß-spectrin (spectrin B2) and protein 4.1R (Fig. 1B and C), before being imaged by STED microscopy. The STED images were subsequently processed and defined fluorescence clusters were observed. Using custom-made algorithms, the cluster distribution densities, their sizes in full width at half maximum (FWHM), the nearest neighbor distances, and the spatial relationship between the two targets were calculated (Fig. 1C) (see Material and Methods).

**Fig. 1.**
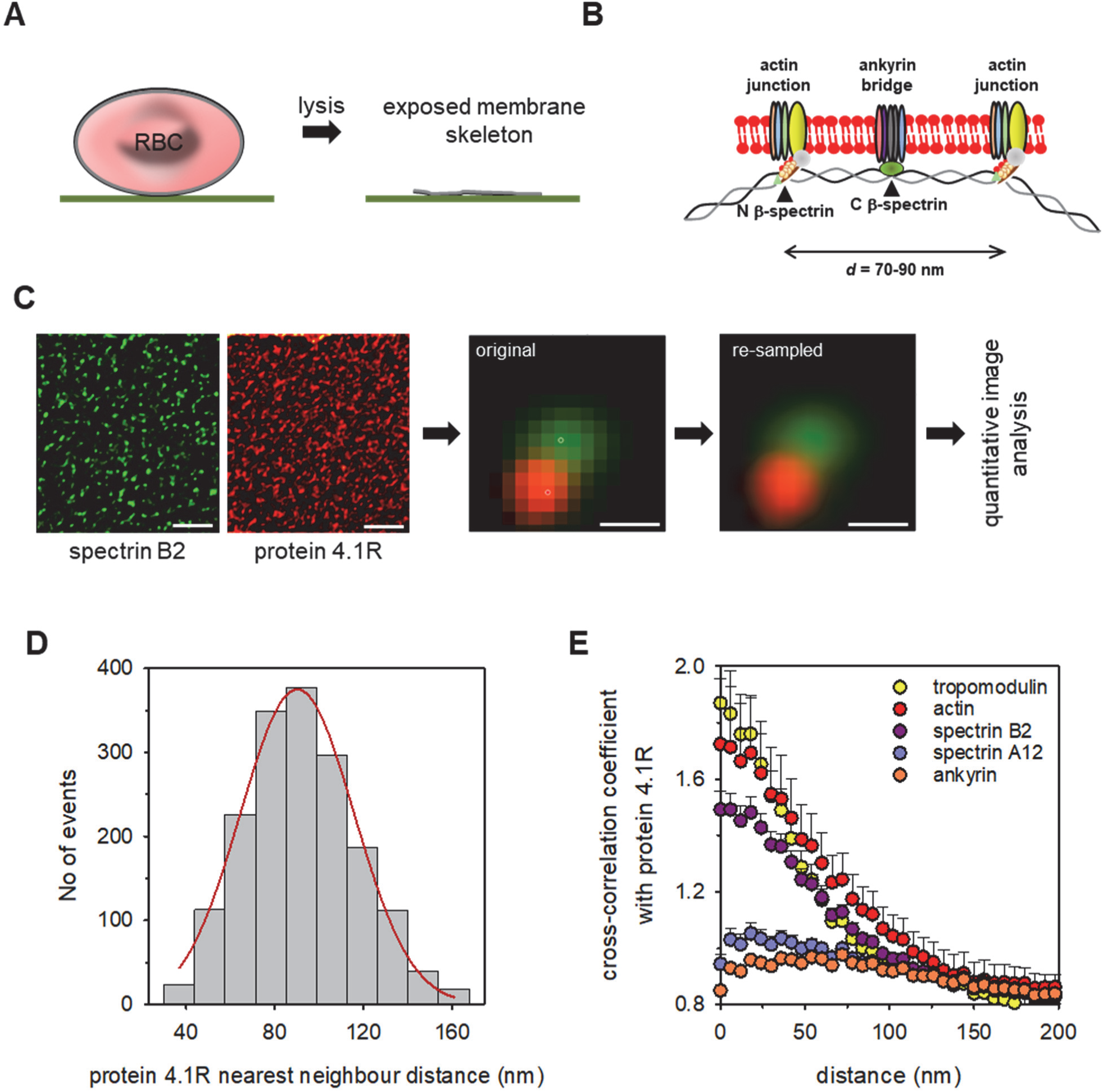
Workflow of STED imaging and method validation. (**A**) Cartoon depicting the preparation of exposed membranes by hypotonic shock of erythrocytes immobilized on a glass slide. RBC, red blood cell. (**B**) Current model of the spectrin/actin network of erythrocytes (see main text for further detail). (**C**) Separate two-color STED images of uninfected erythrocytes stained with an antibody against the N-terminus (acting binding domain) of ß-spectrin (spectrin B2, left panel) and protein 4.1R (right panel). Scale bar, 0.5 µm. Middle panel, zoom-in of overlaid STED images showing both fluorescence clusters. Right panel, same image but re-sampled. Scale bar, 50 nm. (**D**) Distribution of nearest neighbor distance of protein 4.1R. The histogram was fitted using a Gaussian function (red line). N=1471; n=45. (**E**) Calculated two-dimensional cross-correlations between protein 4.1R and tropomodulin (yellow; n=28), actin (red; n=13), N-terminus of ß-spectrin (spectrin B2, purple; n=36), C-terminus of ß-spectrin (spectrin A12, medium purple; n=18) and ankyrin (orange; n=29). The means ± SEM of n cells from at least three different donors are shown. Representative STED images underpinning the cross-correlations are depicted in fig. S3.

We first validated our imaging set-up by comparing the size of the actin and N-terminal ß-spectrin clusters with the known dimension of the underlying molecular structures in the uninfected erythrocyte. The size of the underlying structure is largely defined by the actin protofilaments, which are ∼37 nm long and to which groups of up to six spectrin tetramers attach via the N-terminal domain of ß-spectrin (Lux, 2016). The measured mean of the actin cluster size in FWHM was 45 ± 7 nm (mean ± SD; number of determinations, N=357; number of independent cells analyzed, n=23) and that of the N-termini of ß-spectrin was 48 ± 8 nm (N=2220, n=61) (fig. S1). These values are slightly larger than the reported physical dimension of the protofilament and might be explained by the sideway positions of the spectrin binding sites and the additional sizes of the primary and secondary antibody trees to detect the two targets.

As a second validation step, we determined the nearest neighbor distance for selected targets and compared these results with the literature values. The length of the spectrin filaments defines the distance between individual components of the membrane skeleton. Spectrin filaments in erythrocytes have measured lengths of ∼ 50 to 100 nm and in the fully stretched conformation of ∼ 200 nm (Lux, 2016; Pan et al., 2018). The average nearest neighbor distances of the targets, protein 4.1R, N-terminus of ß-spectrin, C-terminus of ß-spectrin (center of spectrin filaments), and ankyrin, followed a Gaussian distribution in each case and were on average 92 ± 25 nm (N=1741, n=45), 111 ± 34 nm (mean ± SD; N=2021, n=45), 112 ± 31 nm (N=3131, n=32), and 110 ± 30 nm (N=2884, n=45), respectively (Fig. 1D and fig. S2). These values indicate a slightly stretched membrane skeleton in the exposed membranes.

To measure the distance between different fluorophores, we used two-dimensional pairwise cross-correlation analysis between the signals recorded in the two different fluorescent channels from the same sample. The resulting coefficient is expected to be, at zero intermolecular distance, >1 for two co-localizing targets, <1 for two excluding targets, and ∼1 for two independent, randomly distributed targets. Cross-correlation between protein 4.1R and other components of the actin junctional complex, including tropomodulin, actin, and the N-terminal domain of ß-spectrin (spectrin B2), revealed maximal values > 1.5 at an intermolecular distance of 0 to 6 nm (the first binning interval in the cross-correlation analysis, see Material and Methods for details), which quickly declined to ∼1 within ∼ 100 nm (Fig. 1E and fig. S3). In comparison, cross-correlations between protein 4.1R and components of the ankyrin bridge, including ankyrin and the C-terminal domain of ß-spectrin (spectrin A12), revealed minimal values at zero intermolecular distances, which rose to a value of ∼1 within ∼30 nm (Fig. 1E and fig. S3). These results agree with the known molecular architecture of the membrane skeleton, with protein 4.1 stabilizing the connection between the N-termini of ß-spectrin and the actin protofilaments at the vertices of the cytoskeletal meshwork and tropomodulin and adducin capping the pointed and barbed ends of actin filaments, respectively (Lux, 2016). Ankyrin and the C-termini of ß-spectrin localize to the edges of the meshwork at a distance of 30 to 35 nm from the actin junctional complex in the native skeleton (Lux, 2016; Pan et al., 2018) (Figure 1B). Thus, the imaging protocol reveals the established architecture of the spectrin/actin network and appears to be suitable to investigate parasite induced changes of it. Further details on the spatial resolution of the experimental set-up are provided in fig. S4.

### Association of KAHRP with actin but not ankyrin in trophozoites

Fig. 2A depicts representative STED images of exposed membranes prepared from *P. falciparum*-infected erythrocyte at the trophozoite stage (28 – 36 hours post invasion, when mature knobs are already formed), stained with either one of two different antibodies against KAHRP: the mouse monoclonal antibody mAb18.2 (epitope within residues 282 to 362) or the rabbit peptide antibody pAb (epitope within residues 288 to 302) (Kilejian et al., 1991b; Taylor et al., 1987; Watermeyer et al., 2016). Dispersed, homogenous signal clusters were visible in each case, with sizes in FWHM of 55 ± 11 nm (N=2800; n=267) and distribution densities of 22 ± 4 µm^-2^ (n=62) (Fig. 2A and fig. S5). We further explored 3-D STED and again observed homogeneous KAHRP cluster signals (fig. S6A). Comparable results were also obtained when KAHRP stained exposed membranes were imaged by direct stochastic optical reconstruction microscopy (dSTORM) (FWHM of 46 ± 12 nm (N=3383; n=14); distribution density of 22 ± 11 µm^-2^ (n=14)) (Fig. 2A and fig. S5). The appearance of the KAHRP clusters was also unaffected by whether or not specimens were fixed with paraformaldehyde (fig. S6B). In contrast, adding even minute amounts of glutaraldehyde as fixative substantially reduced staining efficiency and the overall quality of the super-resolution images (fig. S6B and C), consistent with previous reports (Mehnert et al., 2019).

**Fig. 2.**
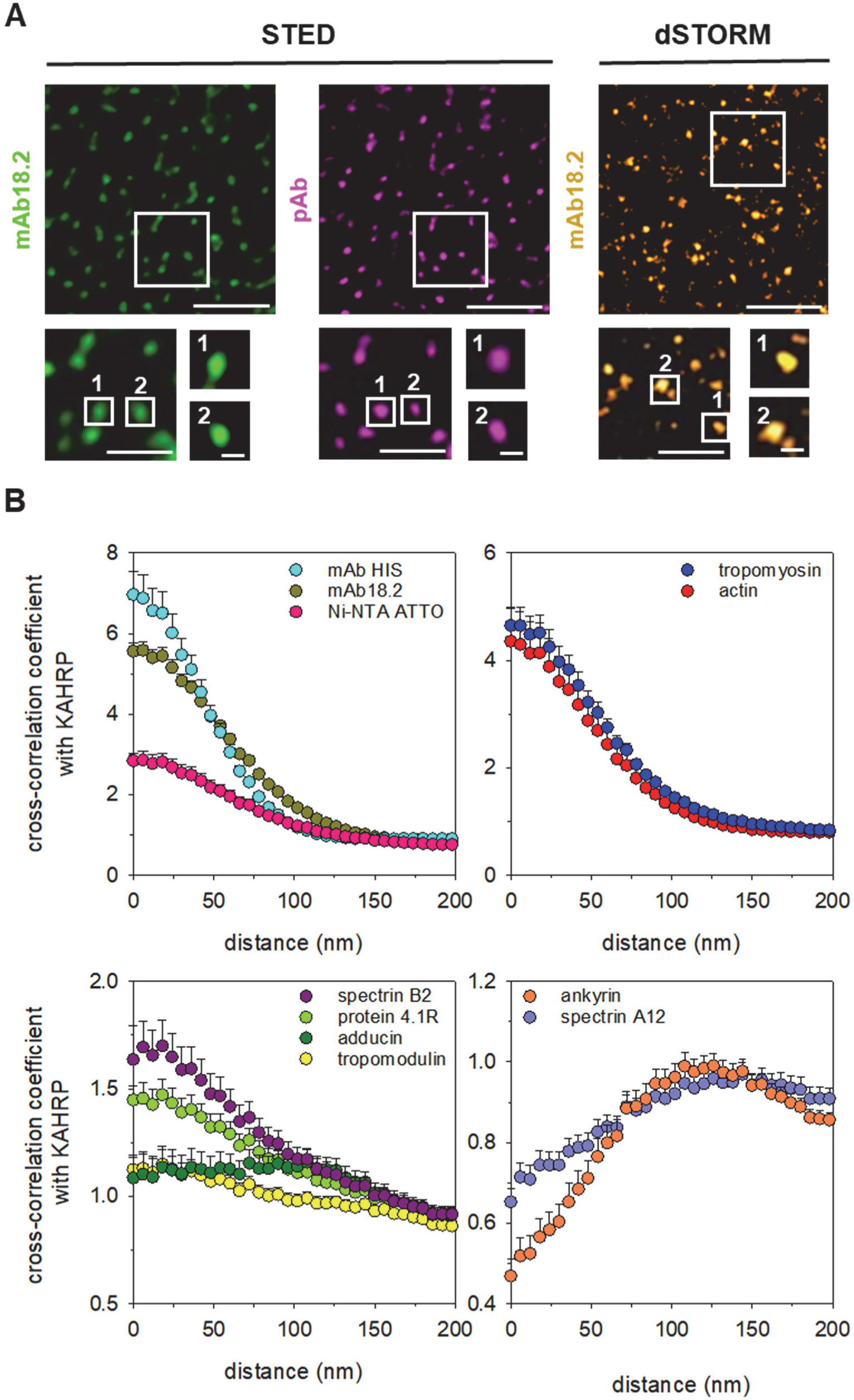
Differential co-localization of KAHRP with components of the trophozoite membrane skeleton. (**A**) STED image of exposed membranes prepared from trophozoites stained with the monoclonal anti-KAHRP antibody mAB18.2 (left panel) or the anti-KAHRP peptide antiserum pAB (middle panel). Right panel, dSTORM image of exposed membranes prepared from trophozoites stained with the monoclonal anti-KAHRP antibody mAB18.2. Zoom-ins of boxed areas are shown below. Scale bars, upper panels, 1 µm; lower left panels, 0.5 µm; lower right panels, 50 nm. (**B**) Calculated two-dimensional cross-correlations between KAHRP (using pAB) and: upper left panel: target of monoclonal anti-His antiserum (cyan; n=15); KAHRP (using mAB18.2; dark olive green; n=99), and target of Ni^2+^-NAT-ATTO 647 nanoprobe (magenta; n=15); upper right panel: tropomyosin (blue; n=36), actin (red; n=58); lower left panel: N-terminus of ß-spectrin (spectrin B2, purple; n=46), protein 4.1R (light green, n=15), adducin (dark green; n=67), tropomodulin (yellow; n=84); lower right panel: C-terminus of ß-spectrin (spectrin A12, medium purple; n=54) and ankyrin (orange; n=39). The means ± SEM of n cells from at least three different donors are shown. STED images underpinning the cross-correlations are depicted in fig. S7.

Two-color STED, using the two KAHRP antibodies, revealed the expected co-localization of the corresponding signals and, accordingly, a high coefficient of 5.9 at 0-6 nm distance in the cross-correlation analysis (Fig. 2B and fig. S7). Co-localization at 0-6 nm distance was also observed between KAHRP and tropomyosin and actin (cross-correlation coefficient >4) and, with coefficients between 1.5 and 1.7, between KAHRP and the N-terminus of ß-spectrin and protein 4.1R (Fig. 2B and fig. S7). The two actin associated proteins, adducin and tropomodulin, displayed an independent spatial correlation with KAHRP, with coefficients of ∼1 (Fig. 2B and fig. S7). In comparison, KAHRP was found to be anti-correlated with the C-terminus of ß-spectrin and ankyrin. In these cases, the cross-correlation coefficients reached a minimum at 0 nm distance before it approached a value of ∼1 at larger distances (Fig. 2B and fig. S7). The strong cross-correlation between KAHRP and actin is consistent with previous cryo-electron tomographic analysis showing long actin filaments connecting the knobs with Maurer’s clefts in trophozoites (Cyrklaff et al., 2012; Cyrklaff et al., 2011; Cyrklaff et al., 2016). Two-color STED imaging further confirmed co-localization of KAHRP with PfEMP1 and PHIS1605w (Cutts et al., 2017; Oberli et al., 2014), with cross-correlation coefficients >2 at 0-6 nm distance, but not with the Maurer’s cleft-associated histidine-rich protein 1 (MAHRP1) (Warncke et al., 2016) (fig. S8).

### Repositioning of KAHRP during parasite development

Previous in vitro studies have revealed that KAHRP is a multi-domain protein with disordered regions that can interact with actin and spectrin and also with ankyrin (Cutts et al., 2017; Warncke and Beck, 2019). We therefore wondered whether the observed strong correlation of KAHRP with actin and the N-terminus of ß-spectrin and the anti-correlation with ankyrin and the C-terminus of ß-spectrin in trophozoites represents only a snapshot of a more dynamic process. To address this hypothesis, we repeated the analysis, but this time using highly synchronized parasites at 12 ± 2, 16 ± 2, and 20 ± 2 hours post invasion. Cross-correlation analyses of two-color STED images showed maximal values at 0-6 nm distances between KAHRP and targets of both the actin junctional complex (N-terminus of ß-spectrin) and the ankyrin bridge (C-terminus of ß-spectrin and ankyrin) at the earliest time point investigated (Fig. 3A and fig. S9A). Comparable results were found for KAHRP at 16 ± 2 hours post invasion (Fig. 3B and fig. S9B). However, 20 ± 2 hours post invasion, the spatial association with KAHRP and ankyrin dramatically changed and the co-localization became statistically independent, with a constant cross-correlation coefficient of ∼1 (Fig. 3C and fig. S9C). In comparison, the high cross-correlation at 0-6 nm distance with KAHRP and the N-terminus of ß-spectrin remained. Plotting the 0-6 nm distance cross-correlation coefficient as a function of time post invasion, highlights the dynamic character of the spatial association between KAHRP and ankyrin - which changed from a strong positive correlation (coefficient >1), to a neutral one (coefficient ∼1) to an anti-correlation (coefficient <1) (Fig. 3D). The temporal nature of the spatial association between KAHRP and ankyrin is further evident when analyzing the ratio of the KAHRP-ß-spectrin N-terminus cross-correlation coefficients at 0-6 nm distance to the corresponding KAHRP-ankyrin values (Fig. 3D). Concomitant with the declining association of KAHRP with ankyrin, the KAHRP cluster sizes in FWHM significantly increased from 51 ± 11 nm (N=2071; n=107) to 55 ± 1 nm (N=2800; n=267) (p<0.001; two-tailed t-test), as the parasite matured (fig. S10). Similarly, the KAHRP cluster density rose from 11 ± 6 µm^-2^ (n=92) to 22 ± 4 µm^-2^ (n=62) (p<0.001; two-tailed t-test) (fig. S10), consistent with previous reports (Looker et al., 2019; Quadt et al., 2012).

**Fig. 3.**
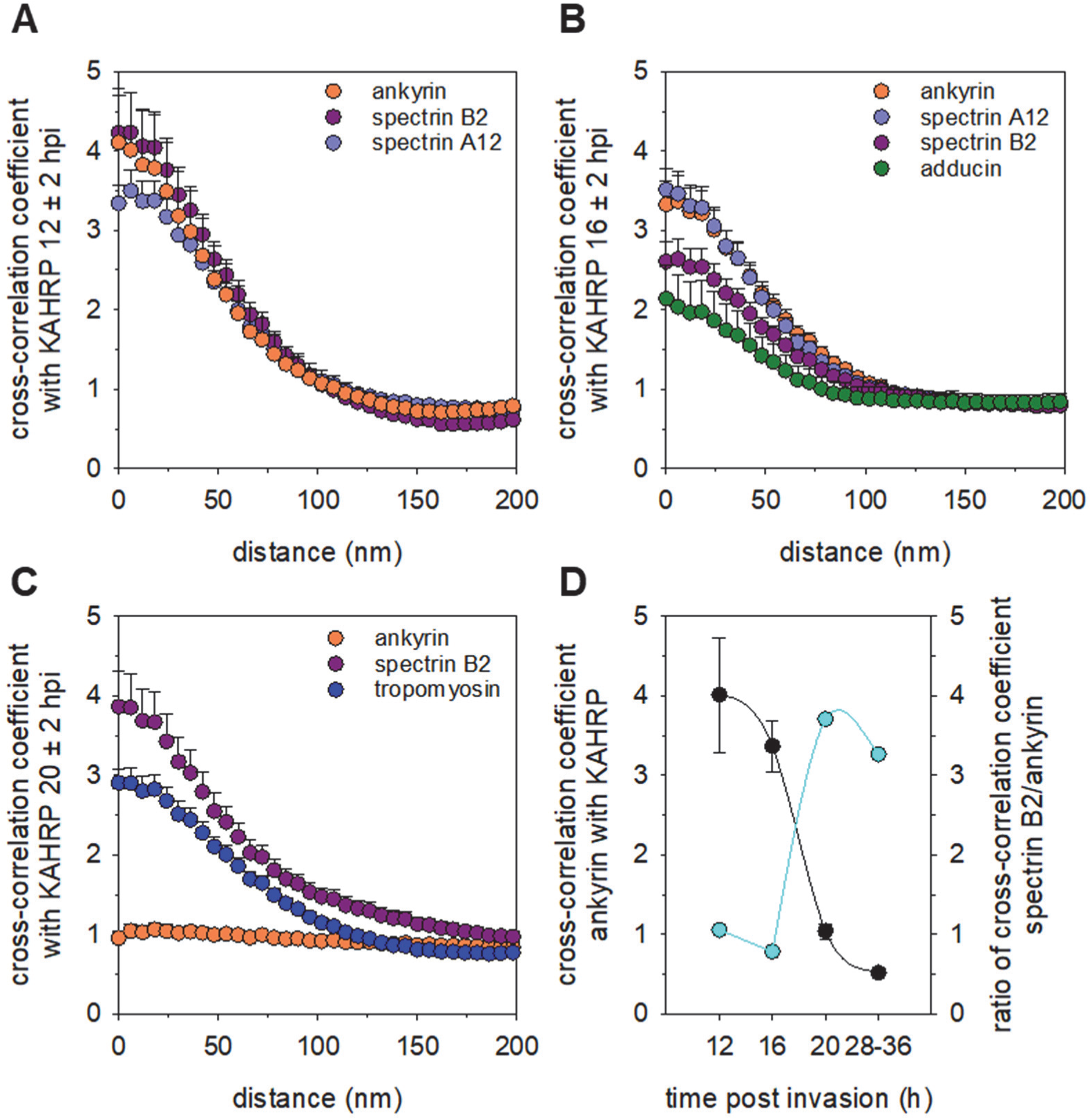
Dynamic co-localization of KAHRP with membrane skeletal components during early parasite development. (**A**) Exposed membranes were prepared from highly synchronized *P. falciparum* cultures 12 ± 2 hours post invasion and stained with an anti-KAHRP antiserum and antisera against the N-terminus of ß-spectrin (spectrin B2, purple; n=15), the C-terminus of ß-spectrin (spectrin A12, medium purple; n=18) and ankyrin (orange; n=22). The calculated two-dimensional cross-correlations between KAHRP and the membrane skeletal components investigated are shown. The means ± SEM of n cells from at least three different donors are shown. STED images underpinning the cross-correlations are depicted in fig. S9. (**B**) as in (A) but investigating parasites 16 ± 2 hours post invasion. C-terminus of ß-spectrin (spectrin A12, medium purple; n=28), N-terminus of ß-spectrin (spectrin B2, purple; n=31), ankyrin (orange; n=28) and adducin (dark green; n=19). (**C**) as in (A) but investigating parasites 20 ± 2 hours post invasion. N-terminus of ß-spectrin (spectrin B2, purple; n=39), ankyrin (orange; n=21), and tropomyosin (blue; n=27). (**D**) Temporal colocalization of KAHRP with ankyrin. The cross-correlation coefficients at 0-6 nm distance between ankyrin and KAHRP were analyzed as a function of the time post invasion (black data points). The mean ± SEM is shown. The ratios of the KAHRP-spectrin B2 cross-correlation coefficients at 0-6 nm distance to the corresponding KAHRP-ankyrin values are shown as a function of parasite development (cyan data points).

### 3D structure of knob spirals

To better understand the structural scaffold underpinning knobs, we applied cryo-electron tomography (cryo-ET) and subtomogram averaging to ghosts prepared from FCR3 infected erythrocytes, preserved by rapid freezing in vitreous ice (fig. 9A and B). The tomograms revealed the erythrocyte plasma membrane and knobs in top and side views (Figure 4A). The most prominent feature was the spiral scaffold underlying the knobs (Figure 4A and fig. S11C), consistent with previous reports (Watermeyer et al., 2016), but here resolved to higher resolution. The spirals had at least 4 to 5 turns, occasionally six or more (Fig. 4B and fig. S11C), and average basal diameters between 55 and 64 nm, corresponding to the FWHM of the KAHRP signal in STED images of mature knobs (fig. S5A). Some spirals were smaller and were only 27-34 nm in diameter, which corresponded to one or two turns. These smaller spirals might represent intermediate or pre-mature structures (Looker et al., 2019).

**Fig. 4.**
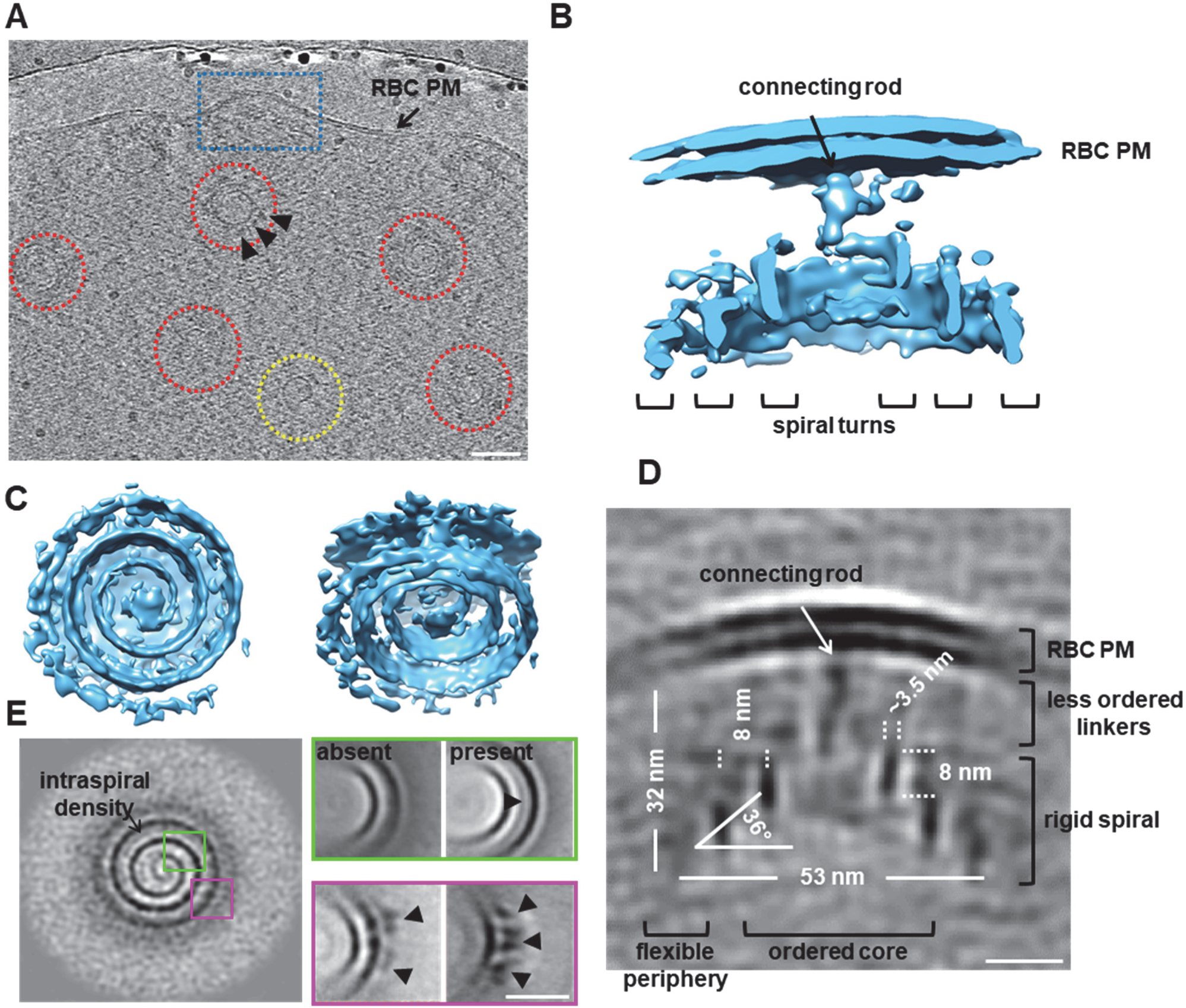
Cryo-ET of the knob spiral. (**A**) A section through a cryo-electron tomogram showing knobs and the underlying spiral in top view (red) and side view (blue). A presumably premature spiral is highlighted in yellow. Scale bar, 50 nm. (**B**) Volume rendered presentation of density map of the knob spiral in cross-sectional view. The plasma membrane of the red blood cell (RBC PM), the connecting rod between spiral and plasma membrane, and the spiral turns are indicated. (**C**) Bottom and skew view of the knob spiral. (**D**) Slice though the average map of the knob with dimensions. Scale bar, 20 nm. (**E**) Averaged tomogram of the knob spiral in top view highlighting intra-spiral densities between the 2^nd^ and 3^rd^ turn in some but not all spirals (green box) and crown-like densities at the outer turn (magenta box). Scale bar, 20 nm.

We next applied subtomogram averaging to the 489 knobs identified in 43 tomograms, yielding a 35 Å resolution density map (332 particles in the final map) (Fig. 4B and fig. 9D and E). The resulting map demonstrates the ordered knob core with an underlying four-turn spiral attached to the membrane of the erythrocyte with a rod-like connector (Figs. 4B and C). The spiral is ∼32 nm in height and ∼53 nm in basal diameter (between centers of mass), and the spacing between each turn is ∼8.0 nm (Fig. 4D). The total spiral blade is 760 nm in length, 3.5 nm in width (limited by the resolution of 3.5 nm) and 8.0 nm in height. The spiral rose at an angle of ∼36° and had a rod-like structure at its tip that extended towards the inner leaflet of the erythrocyte plasma membrane (Figs. 4B and D). There were further multiple less-ordered, stick-like densities in between the erythrocyte plasma membrane and the spiral, which might link the spiral to the plasma membrane (Figs. 4B and D) (Watermeyer et al., 2016). Although we observed spirals with 5 or more turns, the additional turns could not be resolved in our map because the peripheral regions were less ordered (fig. S11) and, thus, were averaged out.

We next performed local classification of the densities of the spiral turns using the subboxing functionality of Dynamo (Fig. 4E, Materials and Methods). Subboxing allows multiple smaller parts to be extracted from subtomograms and to be aligned and averaged locally, as opposed to a global alignment of the entire subtomogram. The subboxing classification revealed intra-spiral densities between the 2^nd^ and the 3^rd^ turn in some but not all knob spirals and crown-shaped densities at the outer turn (Fig. 4E), the latter were also apparent in tomograms at the periphery of knobs (Fig. 4A). The observed intra-spiral densities (Fig. 4A) (Watermeyer et al., 2016) might correspond to associated proteins or protein complexes.

### KAHRP associates with the spiral scaffold

Previous studies have suggested that KAHRP forms an electron-dense coat around the spiral scaffold (Looker et al., 2019; Watermeyer et al., 2016). This conclusion was drawn from cryo-ET analysis of schizont skeletons labeled with a KAHRP antibody and a 10 nm gold conjugated secondary antibody. However, the localization precision of conventional indirect immuno-labeling is limited by the combined sizes of the primary and secondary antibody of up to 30 nm (Vicidomini et al., 2018). To better resolve the location of untagged KAHRP in relation to the spiral scaffold, we looked for alternative labeling techniques and realized that KAHRP contains three stretches of 6 or more histidines within its N-terminal domain, which might be targeted by Ni^2+^-NAT nanoprobes. Ni^2+^-NAT nanoprobes are ∼10 fold smaller than antibody trees and can deliver a gold particle or a fluorescent dye within 1.5 nm of the target (Reddy et al., 2005). To explore the possibility of labeling KAHRP with nanoprobes, we initially tested, by two color STED microscopy, whether the histidine stretches are accessible using an anti-His monoclonal antibody as indicated by the corresponding signal co-localizing with that of an anti-KAHRP antibody in a cross-correlation analysis. The high cross-correlation coefficient at zero nm distance of 7 indeed suggested that the His stretches of KAHRP are targeted by the anti-His monoclonal antibody (Fig. 2B and fig. S7A). Having established accessibility of the His stretches, we next used a Ni^2+^-NAT-ATTO nanoprobe, and again observed co-localization and a high cross-correlation coefficient at zero nm distance with the anti-KAHRP antibody signal (Fig. 2B and fig. S7A).

We next investigated the localization of KAHRP, using immuno-electron tomography. To this end, ghosts from infected erythrocytes (trophozoites) were permeabilized with 0.2% NP40 substitute, labeled with a Ni^2+^-NTA-5 nm gold nanoprobe (Fig. 5A), and embedded in a mixture of uranyl acetate and methylcellulose for electron microscopy (EM) imaging. Fig. 5B depicts an EM projection and a selected knob structure in consecutive tomographic z-sections, showing several spirals in top view with specific gold labeling in close proximity. Importantly, a magnified view of a representative knob shows that the gold particles are placed in the immediate vicinity of the spiral blade and that they closely follow the turns from top to bottom through the z-sections (Fig. 5B). Comparable results were obtained when hosts were not permeabilized or permeabilized with 1% NP40 substitute (fig. S12A and B). To analyze the spatial relationship of the nanoprobe with the spiral we applied stereology methods. To this end, we extracted 27 typical knobs from tomograms and determined the radial distance (r) in relation to the spiral central axis and the z-position in reference to the spiral tip for all associated gold particles (center of mass) (Fig. 5C). A total of 1589 gold particles were mapped. 24% of the gold particles mapped between spiral blades, 21% were slightly offset from the blade towards the interior of the spiral and 55% appeared offset outwards in relation to the spiral blade. We subsequently modeled the blade of the spiral as an Archimedean conical spiral (Fig. 5D), using the parameters determined form the cryo-ET mapping depicted in Fig. 4D. We next simulated the binding of the gold particles to the spiral, with the distance from the center of mass of the gold particle to the spiral varying from 0 to 10 nm. The simulated binding profiles were subsequently compared with the experimental data using a least-squares regression analysis. A distance of 6 nm from the gold particle center of mass to the spiral wall yielded the best fit between simulated and experimental data (Fig. 5E and F). This value is in good agreement with the distance between the center of mass of the gold particle and the His-tag. The agreement in the inclination angle of the simulated and experimental binding profile (Fig. 5F) further supports the hypothesis that the gold particles indeed associate with the spiral surface. Overall, the labelling data suggest that KAHRP is an important component of the knob spirals.

**Fig. 5.**
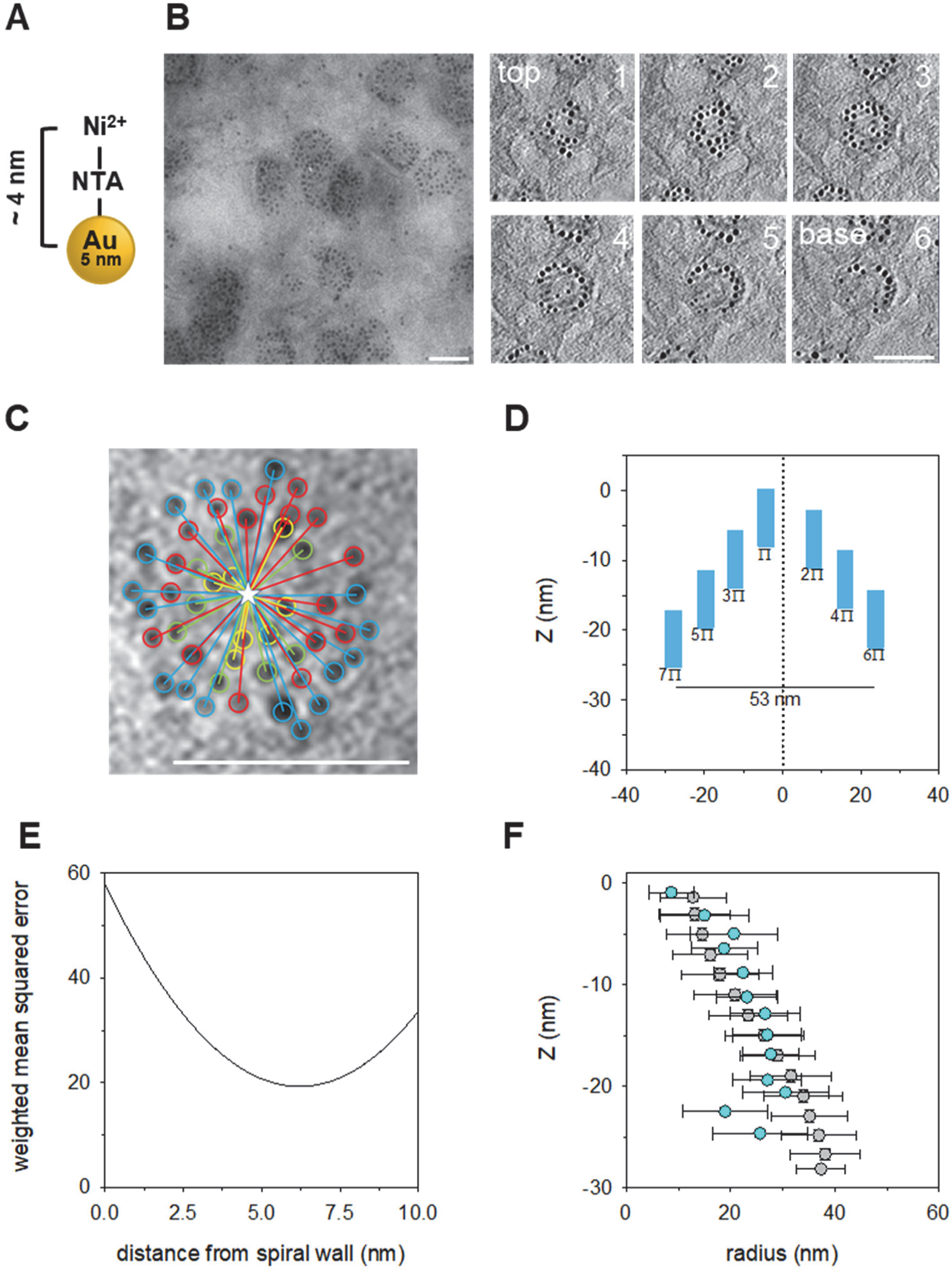
Immuno-EM labeling of the knob spiral with a Ni^2+^-NTA 5 nm gold nanoprobe. (**A**) Cartoon depicting the Ni^2+^-NTA 5 nm gold nanoprobe. The distance from the center of mass of the gold particle to the target is ∼4 nm. (**B**) Left: a projection image showing a segment of a methyl cellulose embedded trophozoite membrane ghost with gold labeling over knobs. Right: consecutive z-sections of an EM-tomogram of a spiral scaffold labeled with the Ni^2+^-NTA 5 nm gold nanoprobe. The sections are numbered in rising order from the top of the spiral to the base. Scale bars, 50 nm. (**C**) Projection of an EM-tomogram of a representative spiral decorated with the Ni^2+^-NTA gold nanoprobe. To determine the coordinates of the gold particles, the spiral top was marked by a yellow star and the radial distance of the gold particle (center of mass) to the spiral vertical center was measured, as was the z-position of the gold particle. The colors indicate gold particles located in different z-sections. Scale bar, 50 nm. (**D**) Simulation of the knob spiral as an Archimedean conical spiral with 4 turns and a basal diameter of 53 nm. (**E**) The binding profile of the gold particles in reference to the spiral blade was simulated at distances from 0 to 10 nm. The simulated data were subsequently compared with the experimental data using a least-squares regression analysis. The weighted mean squared error between the simulated and experimental data is plotted as the distance from the gold particle (center of mass) to the wall of the spiral blade. The minimum of the curve indicates the model with the best fit. (**F**) Overly of the experimental data (cyan) with the best fit model assuming a distance between the wall of the spiral blade and the center of mass of the gold particle of 6 nm (grey). The z-coordinates of the gold particles were plotted as a function of their radial distance from the spiral wall. Error bars indicate the SD.

### KAHRP is a major numeric component of knobs

To quantify the number of KAHRP molecules per knob, we performed quantitative single-molecule localization microscopy (Dietz and Heilemann, 2019; Sanchez et al., 2019). To this end, we inserted one copy of the coding sequence of the photoactivatable fluorescence protein mEOS2 into the 5’ coding region of the genomic *kahrp* gene (corresponding to an insertion between amino acids 207 and 208), using CRISPR/Cas9 genome editing technology (Ghorbal et al., 2014) (Fig. 6A and fig. S12A). Five clonal lines were obtained and the insertion event was confirmed by sequencing of the genomic *kahrp* locus and the *kahrp* transcript. The clone G8 was chosen for further analysis. G8 displayed a typical knobby surface, as shown by scanning electron microscopy (Fig. 6B). However, the mean knob density of 20 ± 6 µm^-2^ was ∼17 % lower than that of the parental strain FCR3 (24 ± 4 µm^-2^) (Fig. 6B). Western analysis confirmed expression of a 25 kDa larger KAHRP protein (fig. S13B) and comparison of a confocal fluorescence image revealing mEOS2 with a STED image depicting KAHRP stained with mAb18.2 showed co-localization of both signals (fig. S13C). dSTORM super-resolution microscopy (Heilemann et al., 2008), using the mAB18.2, revealed dispersed punctate signal clusters in G8 (Fig. 6C), comparable to those seen for the parental line FCR3 (Fig. 2A). The KAHRP clusters were homogenous and 48 ± 26 nm in size in FWHM (N=1708; n=30) (FCR3, 46 ± 30 nm; N=3383; n=14) and had a distribution density of 21 ± 2 cluster µm^-2^ (n=24) (FCR3, 21 ± 11 cluster µm^-2^; n=14) (figs. S5 and S13E). There were no statistical differences between KAHRP cluster size and distribution density between G8 and the parental line FCR3 (two tailed t-test).

**Fig. 6.**
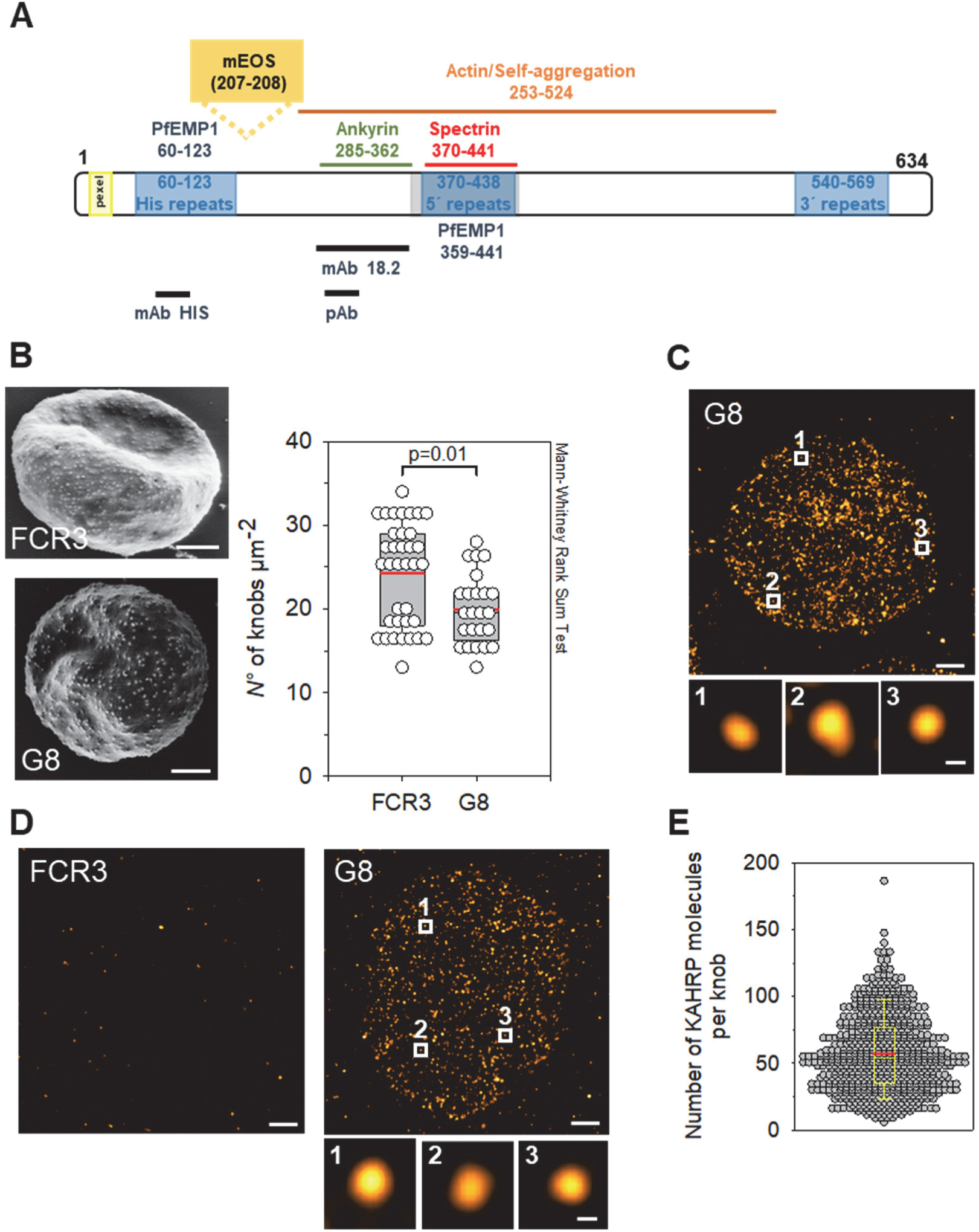
Single-molecule localization microscopy of KAHRP. (**A**) Schematic illustration of KAHRP including structural features and interactions domains. The domains harboring epitopes for the monoclonal antibody mAB18.1, the peptide antiserum and the monoclonal anti-histidine antibody are indicated. mEOS2 was inserted between residues 207 and 208. (**B**) Representative scanning electron microscopic images of erythrocytes infected with the parental line FCR3 or the genetically engineered clone G8 expressing a genomically encoded KAHRP/mEOS2 fusion protein. Scale bar, 1 µm. The knob density per cell was determined and assessed by box plot analyses. Box plots show the individual data points, with the median (black horizontal line), the mean (red horizontal line) and the 25% and 75% quartile ranges being shown. Error bars indicate the 10^th^ and 90^th^ percentile. Statistical significance was assessed using the Mann-Whitney Rank Sum Test. (**C**) Representative dSTORM image of exposed membrane prepared from G8 trophozoites stained with the monoclonal anti-KAHRP antibody mAB18.2. Scale bar, 1 µm. Zoom-ins of boxed areas are shown below. Scale bar 50 nm. (**D**) Representative PALM images of FCR3 and the mutant G8 at the trophozoite stage. The numbered boxes refer to magnified fluorescence clusters presented in the lower panel column. Scale bars, 1 µm in main panel and 50 nm in subpanels. (**E**) Number of KAHRP molecules per knob for G8 at the trophozoite stage, as determined by PALM. A total of 726 clusters from 15 cells were analyzed. A box plot analysis is overlaid over the individual data points.

Fig. 6D shows representative reconstructed PALM images (Betzig et al., 2006) of erythrocytes infected with FCR3 and G8 at the trophozoite stage. Videos were recorded at a frame rate of 10 Hz until no more emission events of mEOS2 were observed (between 150.000 and 200.000 images, 10 Hz), using a total internal reflection fluorescence (TIRF) microscope with an x/y in-plane localization precision ∼14 nm and a light penetration depth of 100 ∼nm. The focal plane was placed on the cell surface and, hence, the knobs. Numerous, punctate fluorescence clusters were observed for G8 (Fig. 6D), with an average size in FWHM of 55 ± 3 nm (N=237, n=13) (fig. S13D). No specific fluorescence signals were seen for age-matched erythrocytes infected with FCR3 (Fig. 6D).

To convert the mEOS2 fluorescence signal into absolute molecule numbers, we considered the following parameters (Hummer et al., 2016; Sanchez et al., 2019): i) an average detection efficiency of mEOS2 of 70%, ii) an average 2.5 detection events per mEOS2 fluorophore, and iii) one mEOS2 copy per KAHRP molecule. Taking these experimentally determined conversion factors into consideration, we obtained 60 ± 30 (mean ± SD) KAHRP molecules per knob (range 10 to 150) (Fig. 6E). This value was based on the analysis of 724 fluorescence clusters from 15 cells, recorded during four sessions, each using blood from a different donor for infection of the G8 line. These results, together with the gold labelling cryo-ET experiments, suggest that KAHRP is present in a surprisingly high amount in knobs.

## Discussion

KAHRP is a major virulence factor during *P. falciparum* infections. Here we have, for the first time, imaged its dynamic localization by combining different quantitative imaging methods, namely optical microscopy with STED, dSTORM and PALM as well as electron tomography combined with labelling and in cryo conditions. To be successful, our approach required the use of specific labeling strategies and the development of adapted algorithms for image analysis. These advances allowed us to map the physical position of KAHRP within the membrane skeleton of infected red blood cells and in relation to the knob spiral in a dynamic manner. Our main findings are that KAHRP dynamically relocalizes during parasite development and eventually forms a major associated component of the spirals underlying the knobs.

KAHRP signals consistently appeared as homogeneous clusters, with a size of ∼50-55 nm, independent of the super-resolution technique used and independent of whether KAHRP was labeled by antibodies or nanoprobes or was tagged with the fluorescent protein mEOS2 (Figs. 2 and 6). Ring- or doughnut-shaped structures with an outer diameter of ∼134 nm and an inner diameter of ∼69 nm, as reported earlier (Looker et al., 2019), were not observed. In fact, such a structure could not have gone unnoticed in our optical setups, which all have resolutions better than 50 nm. Furthermore, gold labelling in electron microscopy showed a uniform distribution of KARHP around the knob spirals. It is unlikely that the organization of KAHRP varies between different strains given the essential nature of KAHRP for knob formation and cytoadhesion in flow (Crabb et al., 1997). A more plausible explanation would seem to be differences in sample preparation affecting morphological preservation and/or access of antibodies to target sites. Particularly, fixation with even minute amounts of glutaraldehyde can detrimentally affect the quality and efficiency of immunofluorescence staining of *P. falciparum* derived specimens (Mehnert et al., 2019) (fig. S6).

KAHRP possesses modular low affinity binding domains that can promote self-aggregation and interactions with parasite factors and components of the membrane skeleton, as shown in in vitro assays (Cutts et al., 2017; Waller et al., 1999; Warncke and Beck, 2019). This includes binding to ankyrin, spectrin, and actin (Cutts et al., 2017; Kilejian et al., 1991a; Oh et al., 2000; Pei et al., 2005; Weng et al., 2014). Current models on the organization of knobs have assumed a static association of KAHRP with these components. Our data suggest a much more dynamic picture. Since our study is time-resolved, we were able to show that the interaction of KAHRP with ankyrin and the C-terminus of ß-spectrin (previously referred to as a ternary KAHRP-spectrin-ankyrin complex (Cutts et al., 2017)) is temporal and occurred only during early ring stage development (Fig. 3), whereas the later trophozoite stage displayed a profound anti-correlation between KAHRP and the two components of the ankyrin bridge (Fig. 2). In comparison, the co-localization of KAHRP with some components of the actin junctional complex persisted throughout intraerythrocytic development, as shown by two-color STED microscopy and pairwise cross-correlation analyses (Figs. 2 and 3).

The temporal co-localization of KAHRP with ankyrin bridge components would suggest a model in which KAHRP reorganizes its own clusters during knob formation or, alternatively, it might reflect a step in the parasite-induced reorganization and disassembly of the spectrin/actin network. Reorganization of KAHRP might be mediated by the changing phosphorylation and/or acetylation pattern of KAHRP during the intra-erythrocytic cycle (Cobbold et al., 2016; Pease et al., 2013), thereby altering the affinity of the protein to components of the membrane skeleton. An initial affinity to multiple membrane skeletal components has the clear advantage of creating a concentration gradient away from the parasite, which allows passive transport by diffusion and does not require yet active transport, e.g., along the long actin filaments that form in later stages. Once accumulated at the red blood cell membrane with sufficient concentration, KAHRP then could reorganizes to its final destination.

Previous models on knob organization have placed KAHRP and the spiral scaffold in the mesh formed by spectrin filaments (Looker et al., 2019), close to the ankyrin bridge (Cutts et al., 2017) or at the actin junction (Oh et al., 1997; Oh et al., 2000). However, these models did not consider the possibility of dynamic rearrangements, involving a transient state at the ankyrin bridge during early ring stage development. Our finding of a spatial correlation between KAHRP and actin and the N-terminus of ß-spectrin in trophozoites suggests that KAHRP eventually assembles at actin junctional complexes (Figs. 2 and 3). While the actin junction may nucleate knob formation, we do not think that mature knobs maintain the anatomy of the actin junction. Previous cryo-ET has revealed that the parasite mines the actin from protofilaments to generate long actin filaments connecting the knobs with the Maurer’s clefts and serving as cables for vesicular trafficking of PfEMP1 and other parasite factors to the red blood cell surface (Cyrklaff et al., 2012; Cyrklaff et al., 2011; Cyrklaff et al., 2016). In addition, the spectrin filaments stretch and the mesh size increases (Shi et al., 2013), which, in turn, contributes to membrane stiffening (Fröhlich et al., 2019; Lai et al., 2015). Consistent with dissembled actin junctions in knobs, we observed an independent spatial behavior between KAHRP and the actin filament capping factors adducin and tropomodulin (Fig. 2B). In comparison, KAHRP strongly correlated with tropomyosin (Fig. 2B), which might suggest that the long actin filaments extending from knobs are stabilized by tropomyosin at least at sites where they are close to, or connected with, KAHRP.

Our study further sheds new light on the number of KAHRP molecules per knob. According to our quantitative PALM experiments, knobs contain on average 60 ± 30 KAHRP molecules (range 10 to 150). This number is more than one order of magnitude higher than previously reported (Looker et al., 2019) and would suggest that KAHRP is a major numeric component of knob protrusions. We considered the possibility that tagging the endogenous, genomically-encoded KAHRP with mEOS2 affected trafficking or function of the resulting fusion protein. However, we do not think that such effects influenced the outcome or the interpretation of our results as the corresponding mutant displayed a knobby phenotype, albeit the knob density was slightly lower according to scanning electron microscopy than that of the parental FCR3 line (Fig. 6B), whereas KAHRP cluster sizes and distribution densities were comparable according to dSTORM imaging (figs. S5 and S13E).

Previous studies have posited the hypothesis of KAHRP coating the spiral scaffold underlying knobs (Looker et al., 2019; Watermeyer et al., 2016). This assumption arose from cryo-ET of schizont skeletons labeled with an anti-KAHRP antibody and a 10 nm gold-conjugated secondary antibody (Watermeyer et al., 2016). However, the combined sizes of the antibody tree and the gold particle of ∼40 nm complicates the interpretation of the results. To improve the localization precision of KAHRP, we used a Ni^2+^-NTA nanoprobe. Ni^2+^-NTA nanoprobes are 10 times smaller than antibody trees and can deliver a gold particle or a fluorescent dye within 1.5 nm of the target (Reddy et al., 2005). Each Ni^2+^ ion coordinates with two histidines and tight binding is achieved when three adjacent Ni^2+^-NTA groups bind to six consecutive histidines (Reddy et al., 2005). KAHRP contains three stretches of 6 or more histidines within its N-terminal domain, making the protein a potential target for Ni^2+^-NAT nanoprobes. We demonstrated feasibility of this approach by showing strong cross-correlation at zero distance and, hence, colocalization of a Ni^2+^-NTA ATTO nanoprobe with an anti-KAHRP antibody in two color STED microscopy of exposed membranes prepared from trophozoites (Fig. 2B). We then showed that a Ni^2+^-NTA 5 nm-gold nanoprobe, in combination with EM-tomography, clearly decorated the spiral blade from top to bottom through the z-sections. The average distance between the center of mass of the gold particle and the spiral wall of 6 nm was in good agreement with the size of the nanoprobe of ∼4 nm (center of mass of the gold particle to the Ni^2+^ ion), as shown by comparing the experimental data with stereological computer simulations (Fig. 5). In addition, the experimentally determined radial distance of the gold particles from spiral central axis matched the inclination angle of the spiral scaffold as determined by cryo-ET (Fig. 5).

The stereological computer simulations were guided by an improved 35 Å resolution density map of the knob complex, which described the geometry of the spiral, defining metric values for its height, width, lengths, basal radius and inclination angle. In this context, it is worth mentioning that the basal radius of the spiral and the KAHRP fluorescence cluster size have comparable values. The 35 Å map of the knob spiral corroborates previous observations (Watermeyer et al., 2016), such as the presence of stick-like densities that seem to anchor the spiral to the lipid bilayer. In addition, our study revealed some additional features, including the intra-spiral densities between the 2^nd^ and 3^rd^ turn, which might stabilize the spiral, and the crown-like extra densities at the periphery of the spiral base, which might be involved in connecting the spiral with filamentous structures.

Although our nanoprobe-based stereological approach cannot replace high-resolution structural information, it nevertheless provided first experimental evidence of KAHRP localizing at, or close to, the spiral blade. On the basis of these findings, we propose that KAHRP is a component or an associated factor of the spiral scaffold and is distributed equally along the spiral length. However, we do not think that the spiral is solely made of KAHRP. Given a molecular weight of 62,682 Da for the processed KAHRP and a volume conversion factor of 1/825 nm^3^ Da^-1^ (Erickson, 2009), this would translate to a predict volume of ∼75 nm^3^ per KAHRP molecule. Dividing the volume of the spiral blade (HxWxL: 8 nm x 3.5 nm x 750 nm) by the volume of a KAHRP molecule would amount to ∼300 KAHRP molecules per average 4- turn spiral if the spiral were made only of KAHRP, considerably larger than the ∼60 KAHRP molecules measured herein. Moreover, as ordered assemblies typically form by association of folded domains, this hypothetical value will rise to ∼700 KAHRP molecules per 4-turn spiral if one considers only the ordered domain of KAHRP from residues 130 to 250. Yet, neither PALM single molecule counting nor EM-tomography using the Ni^2+^-NTA 5 nm-gold nanoprobe support such high numbers. While our study provides evidence of KAHRP being a component or an associated factor of the spiral scaffold further experimental data are needed to verify this hypothesis and to determine other putative constituents. If KAHRP is indeed associated with the spiral blade, then KAHRP might attach the spectrin and actin filaments to the spiral scaffold (Fig. 7). Such a function would be consistent with the binding properties of KAHRP.

**Fig. 7.**
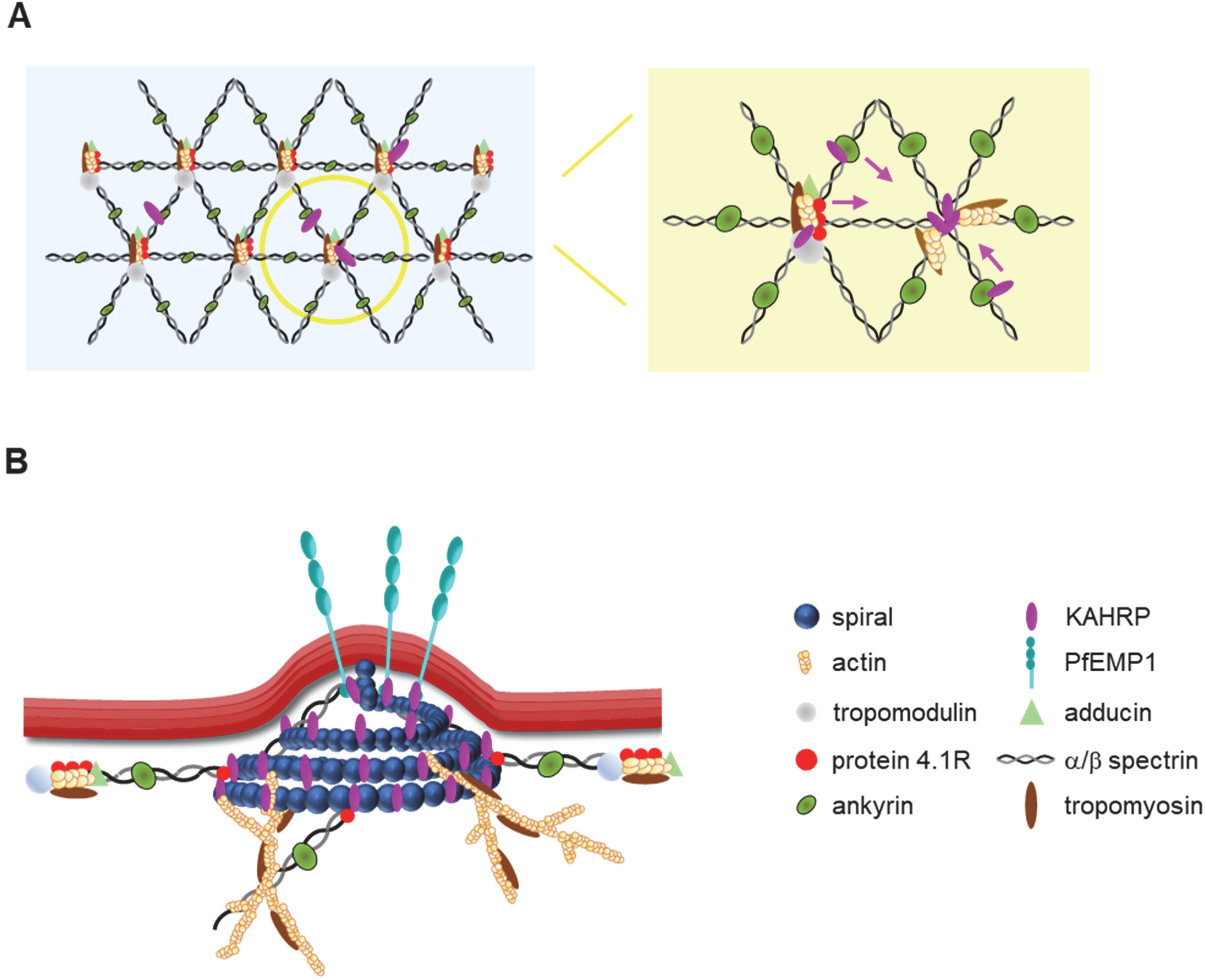
Graphic model depicting the organization of KAHRP. (**A**) Dynamic association of KAHRP with ankyrin bridges. We propose that KAHRP initially binds to both the ankyrin bridge forming a ternary complex with ankyrin and spectrin (Cutts et al., 2017) and to the actin junctional complex during early ring stage development. We further propose that KAHRP then re-positions to the actin junctional complex as the parasite matures. Concomitantly, the knob spiral forms and the actin junctional complexes re-organize. Re-organization of the actin junctional complex would include uncapping and mining the actin protofilaments to form long filaments connecting the knobs with the Maurer’s clefts (Cyrklaff et al., 2011). (*B*) Model depicting KAHRP as an associated factor of the spiral scaffold. According to the model, KAHRP would “glue” additional components to the spiral scaffold. This would include PfEMP1, the spectrin filaments (via a quaternary complex consisting of the N-terminus of ß-spectrin and protein 4.1R), and the long actin filaments. A “glue-like” function would be consistent with the multi-modular binding properties of KAHRP (Warncke and Beck, 2019). The KAHRP symbol indicates one or several KAHRP molecules.

In summary, our findings suggest a dynamic and stage-dependent association of KAHRP with components of the membrane skeleton (Fig. 7A). We propose that KAHRP initially uses multiple binding sites in the red blood cell cytoskeleton to quickly accumulate at the membrane, but later assembles at actin junctional complexes. We further propose that the anatomy of the junctional complex is lost as the knob matures, replacing it by a KAHRP-containing spiral scaffold from which long tropomyosin enforced, uncapped actin filaments extend towards Maurer’s clefts and which stabilizes the actin mined membrane skeleton by bundling the N-terminal ends of ß-spectrin filaments at a cost of an increased mesh size and membrane hardening (Fig. 7A and B).

## Materials and Methods

### Antibodies and nanoprobes

Antisera and nanoprobes used in this study are listed in table S1.

### Oligonucleotides

Oligonucleotides and primers used in this study are listed in table S2.

### Parasite culture

The *P. falciparum* clonal line FCR3 (alias IT2) and the FCR3-derivated clonal mutant G8 were cultured as described (Trager and Jensen, 1976), using fresh HbAA erythrocytes (less than 10 days old after blood donation) resuspended in RPMI 1640 medium supplemented with 5% GlutaMAX (ThemoFisher Scientific) and 5% human serum. The cultures were maintained using the following conditions: 4% haematocrit, <5% parasitemia, 37°C temperature, 5% O_2_, 3% CO_2_ and 96% humidity. Cultures were synchronized using 5% sorbitol (Lambros and Vanderberg, 1979).

### Preparation of exposed membranes

Glass bottom culture dishes (Mattek corporation) were treaded with 2% 3-aminopropyl triethoxysilane (APTES) in 95% ethanol for 10 min, washed with 95% ethanol, and then incubated at 100°C for 15 min before dishes were incubated with 1 mM bis-sulfosuccidimyl suberate in phosphate buffer saline (PBS) at room temperature for 30 min (Shi et al., 2013). Dishes were subsequently washed with PBS and then treated with 0.1 mg ml^-1^ phytohemagglutinin E (PHAE) in PBS, as described (Shi et al., 2013). Dishes were then washed with PBS and blocked with 0.1 M glycine for 15 min, washed once more with PBS and kept at 4°C until further use. Uninfected erythrocytes or magnetic enriched *P. falciparum-*infected erythrocytes were immobilized on the functionalized dishes followed by washes with a hypotonic phosphate buffer (10 mM sodium phosphate pH 8, 10 mM NaCl) followed by washes with water (Dearnley et al., 2016). Exposed membranes were either unfixed, or fixed with 4% paraformaldehyde in PBS or 4% paraformaldehyde and 0.0065% glutaraldehyde in PBS for 15 min, washed with PBS and blocked in PBS containing 3% BSA. Exposed membranes were incubated with primary antibodies overnight at 4°C and with secondary antibodies for 40 min at room temperature. All incubations and washes were performed in 3% BSA in PBS.

### STED microscopy

Super-resolution images were recorded using a STED/RESOLFT microscope (Abberior Instruments, Germany) equipped with 488 nm, 594 nm and 640 nm excitation (Ex) laser sources and 775 nm STED laser lines and an Olympus microscope with a 100× oil immersion objective (UPLSAPO 1.4NA oil, 0.13 mm WD). The STED laser power was adjusted to ∼40%. Fluorescence emitted from Abberior star 580 conjugated secondary antibodies were recorded in the Ex 594 channel (“green images”) and fluorescence emitted from Abberior star red conjugated secondary antibodies were recorded in the Ex 640 channel (“magenta image”). See table S1 for antibodies used in this study. The pixel size was 15 nm and the pixel dwell time was 10 μs. De-convolution of 2D-STED images was performed using the Imspector imaging software (Abberior Instruments GmbH) and the Richardson-Lucy algorithm with default settings and regularization parameter of 1e^-10^. 3D-STED images were de-convoluted using the Huygens Professional 20.04 software (Scientific Volume Imaging B.V.). The differences between sections in z-direction were 150 to 200 nm. In the case of mEOS 2.1 imaging by confocal microscopy, the sample was excited using the 488 nm laser line and the emission was detected using a 500-550 nm bandpass filter.

### Cross-correlation analysis

Two-dimensional cross-correlation between two color channels was computed by calculating the pairwise inter-molecular distance distribution (PDD) from the images (Schnitzbauer et al., 2018). The shortest distance between the points on two images is limited by the size of a pixel (15 nm), which, in turn, limits the PDD at short distances by the size of a pixel. To overcome this limitation, we used linear interpolation (or re-sampling) on the original image to interpolate the signal into smaller grids. This is possible because single fluorescence signals cover multiple pixels. The image interpolation step smoothens the image and improves the PDD (Schnitzbauer et al., 2018). We chose one tenths of a pixel, corresponding to ∼1.5 nm, as the width of the interpolated grid. The resultant PDD was binned into different spatial bins to compute the average PDD at different distances. The bin size was chosen (∼ 6 nm) such that it is larger than one pixel unit of the re-sampled images. The PDD histogram is then radial averaged and normalized by the image area to get the pair cross-correlation (PCC) between two signals for different distances. The PCC is analogous to a cross-correlation function defined for localization points (Sengupta et al., 2011) (see supplementary material and methods).

### Cluster size and distribution

To study cluster statistics, we determine the number of peaks in a given image. The number of peaks divided by the image area defines the cluster density. For each peak, we find the nearest neighbor. The distribution of distances between nearest neighbors is used to estimate the average distance between the clusters. For isolated peaks (with no peaks in 2 pixels or 30 nm distance), we crop a 2 x 2-pixel area and fit the resulting profile with a Gaussian distribution. The full width at half maximum is calculated (FWHM) from the standard deviation (FWHM = 2.355 SD) estimated by the Gaussian fit.

### Cryo-electron tomography, image processing and subtomogram averaging

Highly synchronized parasite cultures at the trophozoite stage were magnet purified and membrane ghost were prepared by incubating the cells in lysis buffer (5 mM Na phosphate pH 7.4, 2 mM MgCl_2_, 1 mM DTT and 1x Halt Protease Inhibitor cocktail (ThermoScientific) for 5 min on ice. After centrifugation at 17,000 x g for 15 min at 4°C, the membranous upper layer was collected and washed several times with lysis buffer. The membrane ghosts were mixed with 10-nm gold fiducial markers in PBS buffer. An aliquot (3 μl) of this mixture was applied to a glow-discharged holey carbon grid (QF R2/2 Cu, QUANTIFOIL), blotted with Whatman® No. 1 filter paper and plunge-frozen in liquid ethane (Vitrobot™ Mark IV, Thermo Fisher Scientific).

Imaging was performed on an FEI Titan Krios microscope fitted with a Gatan Quantum energy filter and a Gatan K2 Summit direct detector operated by Serial-EM software (Schorb et al., 2019). A total of 43 tomographic series was acquired using a dose-symmetric scheme (Hagen et al., 2017), with tilt range ±60°, 3° angular increment and defoci between −5 μm and −6 μm. The acquisition magnification was 42,000 ×, resulting in a calibrated pixel size of 3.39 Å. The electron dose for every untilted image was increased to around 10 e^−^ Å^-2^, and tilt images were recorded as ten-frame movies in counting mode at a dose rate of approximately 0.62 e^−^ Å^−2^ s^−1^ and a total dose per tomogram of around 110 e^−^ Å^−2^.

Motion correction of tilt-series movies was performed using MotionCor2 (Zheng et al., 2017). Tilt-series were aligned on the basis of the gold fiducials using the IMOD package (Kremer et al., 1996). Contrast transfer function (CTF) estimation was performed using defocus values measured by Gctf (Zhang, 2016) for each projection. Tomograms were reconstructed from CTF-corrected, aligned stacks using weighted back-projection in IMOD. Tomograms were further binned 2, and 4 times (hereafter called bin2 and bin4 tomograms), resulting in pixel sizes of 6.78 Å and 13.56 Å, respectively.

Subtomogram averaging was performed using the *Dynamo* package (Castaño-Díez et al., 2012). To define initial subtomogram positions, the center of cubic voxels was manually picked on bin4 tomograms (489 particles), using the Dynamo Catalogue system (Castaño-Díez et al., 2017). Initial alignment was done manually on 90^3^ pixel subtomograms extracted from bin4 tomograms, and this initial averaging was low-pass-filtered to 60 Å as a starting reference for alignment of the bin2 tomograms. Multi-reference alignment and averaging were performed on bin2 particles in 180 voxel boxes. A soft-edged sphere alignment mask was applied throughout and the full dataset was aligned against the core of knobs, revealing prominent spiral-like structures decorating the lipid bilayer. Subsequent iterations of refinement were performed on subtomographs of 360 voxels extracted from unbinned tomograms with constrained refinement of shifts and angular search on statistically independent odd and even datasets. A population of particles with lowest cross-correlation (around 20%) was removed by imposing a cross-correlation threshold. Final converged averages were formed by 332 particles, and the resolution was determined by *dynamo_fsc*. Further alignments were performed using ellipsoid masks focused on two different regions of the spiral structure (middle, and peripheral regions) to generate 4 maps. The subboxes were extracted in 40 positions around the 2^nd^ and 3^rd^ spiral turn of each particle using *dynamo_subboxing_table*. Symmetry was not applied for the consecutive refinements.

### Labeling with a Ni^2+^-NAT-gold nanoprobe

Membrane ghosts were prepared as described above. Ghosts were subsequently extracted with NP-40 substitute (Sigma Aldrich) (10 mM Na phosphate pH 7.4, 2 mM MgCl_2_, 1 mM EDTA, 1 mM DTT, 0.2%, 0.4% or 1.0% NP-40 substitute where indicated) for 5 to 10 min at 4°C before centrifugated at 17,000 x g for 15 min at 4°C. The pellet was collected and re-suspended in a small volume of the NP-40 substitute buffer. 2-3 µl were subsequently placed on a 100 mesh copper grid with pioloform (1.5 % v/w) support film (Peurla et al., 2019) pre-coated with carbon and glow discharged and functionalized with 0.1% poly-L-lysine solution for 5 min. The grids were washed in PBS, incubated for 10 min at room temperature in blocking buffer (1% fish skin gelatin, 20 mM Tris/HCl pH 7.4), and a drop containing the Ni^2+^-NAT-5 nm gold nanoprobe diluted 1:30 in blocking buffer (1% fish skin gelatin, 20 mM Tris/HCl pH 7.4) was placed on the grid and incubated for 30 min at room temperature. Grids were rinsed with washing buffer (20 mM Tris/HCl pH 7.4, 150 mM NaCl, 8 mM imidazole) several times followed by three washes with water. Samples were embedded in a mixture of 1% methylcellulose and 1% uranyl acetate and the grid was air dried overnight.

For electron tomography, the Tecnai F20 transmission electron microscope, which operates with a tungsten field emission gun at 200 kV, was used with a defocus of 1 µm. Tilt-series were recorded within the maximal angles of -60° to +60° with 1-2° increments using SerialEM software for data collection. The digital images were recorded on a CCD-Eagle 4Kx4K camera at a nominal magnification of 25,000 that corresponds to the pixel size of 0.8919 nm. Tilt-series were processed and reconstructed into 3D tomograms using the IMOD software package and fiducial markers (Kremer et al., 1996). The radial distance from the spiral central axis (r) and the distance in z from the spiral top was determined for each associated gold particle. To do so, the center of the spiral was determined and the top was marked with a yellow star. The spiral scaffold structure was subsequently divided into vertical zones along the z-axis, with one z-interval consisting of 5 layers (equivalent to 5 pixels or 4.459 nm in thickness).

### Stereological computer simulation

We simulated the knob spiral as an Archimedean conical spiral. The equations for the spiral are defined by two parameters: the slope of the spiral surface **m** and the radius **r**. The slope is estimated from the y-z cross-section of an average tomogram of the knob spiral. The radius **r** is computed from the width of the 4th turn of the spiral (corresponding to ϕ=5π on the left side and 6π on the right side). The 3D coordinate of points on the spiral is described by x=r ϕ cos(ϕ), y=r ϕ sin(ϕ), z=- m r ϕ. The spiral surface is defined by two contours; an upper contour is defined by x,y,z, and a lower contour is defined by x,y,z-z_thick_, where z_thick_ (= 8 nm) is the height of the spiral strip. To assign particles to the spiral, we distributed them using randomly distributed ϕ (between π to 8π) and z values (between the upper and lower contour). Next, we added Gaussian noise around x,y values with variance similar to the experimentally observed dataset. The binding profile was then plotted by binning the z-coordinates in 2 nm bins. For each bin, the mean and SD of radial distance from the spiral center line and z-coordinate values were estimated. To fit our model, we computed the weighted mean squared error between the experimental data and model predictions (for uniform binding on the spiral surface) for different radial distances (0 to 10 nm) from the center of mass of the particles to the spiral wall. A separation distance of ∼6 nm yielded the best fit between simulated and experimental data. This separation distance is in the range of the distance between the center of mass of the gold particle and the His-tag (Reddy et al., 2005).

### Generation of a *kahrp/mEOS2* mutant

For the generation of the KAHRP-mEOS2.1 mutants, the entire *kahrp* gene was amplified by PCR using FCR3 gDNA and cloned into the pL6-B vector (Ghorbal et al., 2014). A DNA fragment between the AflII and AleI restriction sites (amino acid 208 to 308 of KAHRP) was re-codonized (GeneArt, ThemoFisher Scientific). The fragment was then used to replace the corresponding sequence in the *kahrp* coding sequence cloned in the pL6-B vector, using In Phusion (Takara Bio). Afterwards, the re-codonized mEos2.1 coding sequence was inserted into the AflII site by In Phusion. The guide RNAs were cloned into the BtgZ1 site of pL6-B. The guide RNAs were designed such that they would target the region between the AflII and AleI restriction sites of the endogenous *kahrp* gene. For this reason, this region was replaced by a re- codonized version in the transfection vector. The final transfection vector was verified by sequencing analysis. Transfections were performed under standard conditions, using 75 µg each of the plasmid and the Cas9-expressing vector (Ghorbal et al., 2014). Transfected parasites were selected on 1.5 µM DMSI and 5 nM of WR99210. Integration events were verified by PCR of genomic DNA and sequencing analysis. Clones were obtained by limiting dilution and integration processes were again confirmed by PCR and sequencing of the resulting PCR fragments and the corresponding mRNA. Primers used for cloning and sequencing are listed in table S2.

### Single-molecule localization microscopy

LabTek chambers (ThermoFisher Scientific) were covered with 0.1 mg ml^−1^ concanavalin A for 60 min, washed with water and PBS before magnet purified infected erythrocytes were allow to settle on them for 10 min. After washing with PBS, cells were fixed with 4% paraformaldehyde in PBS for 10 min. Paraformaldehyde was removed and cells were washed and kept in PBS at 4 °C until imaging. All buffers were sterile filtrated, using 0.45 µm membrane filters. Samples were imaged within 48 h after preparation. Single-molecule localization microscopy (SMLM) (Sauer and Heilemann, 2017) was performed on a home-built microscope in total internal reflection fluorescence (TIRF) illumination mode (Karathanasis et al., 2020). In brief, an inverted microscope (Olympus IX71) with a 100x oil immersion objective (PLAPO 100x TIRFM, NA ≥ 1.45, Olympus) is equipped with three laser modules (LBX-405-50-CSB-PP, Oxxius; Sapphire 568 LP, Coherent; LBX-638-180, Oxxius). The fluorescence signal was filtered with a bandpass filter (BrightLine HC 590/20, AHF) and collected using an EMCCD camera (iXon Ultra X-10971, Andor). mEOS2 was photoconverted with 405 nm (0-8 mW cm^-^ ²) and excited with 568 nm (0.26 kWcm^-^²), using an EMCCD integration time of 100 ms, a pre-amplifier gain of 1 and an electron multiplying gain of 200. Quantitative PALM experiments were recorded until no more emission events were observed for mEOS2. SMLM images were reconstituted using the localize software of Picasso (Schnitzbauer et al., 2017) using a Min.Net gradient of 3000 and a filter for the width of point spread functions set to a range of 0.4 - 1.6. Single-molecule emission events that occurred in consecutive frames were linked to one event. Single protein clusters were selected and the number of mEOS2 blinking events per cluster determined. To determine the number of KAHRP proteins per protein cluster, the average number of detection events of mEOS2 was calibrated with single mEOS2 proteins attached to a poly-lysine surface (Fricke et al., 2015). Previous studies have shown that the blinking parameters of poly-lysine bound mEOS2 molecules is comparable to those seen for mEOS2 fusion proteins in cells, likely explained by the barrel structure of the fluorescent proteins protecting the chromophore from the environment (Durisic et al., 2014; Fricke et al., 2015; Krüger et al., 2017; Lee et al., 2012).

dSTORM imaging (Heilemann et al., 2008) was conducted by photoswitching Alexa Fluor 647 with 640 nm (1 kW/cm²) using PBS with 100 mM MEA, protocatechuic acid (PCA) and protocatechuate-3,4-dioxygenase (PCD) in pH 7.8 adjusted with 1 M NaOH. The fluorescence signal was filtered with a bandpass filter (ET 700/75, AHF). dSTORM experiments were performed for 20,000 frames with an integration time of 30 ms, a pre-amplifier gain of 1 and an electron multiplying gain of 200. Images were reconstructed using rapidSTORM (Wolter et al., 2010) with a threshold of 278 photons. Mean values of cluster size and cluster per µm² were determined by DBSCAN (Ram et al., 2010) analysis implemented in LAMA (Malkusch and Heilemann, 2016), using a radius of 30 nm and a minimal number of localizations of 5 per cluster as threshold parameter. The localization precision of dSTORM experiments is ∼12 nm, as determined by nearest-neighbor analyses (Endesfelder et al., 2014).

### Statistical analysis

Data are presented as following: FWHM (nm), mean ± SD; cluster density (µm^-2^), mean ± SD; nearest neighbor distance (nm), mean ± SD; number of KAHRP molecules per knob, mean ± SD; and cross-correlation coefficient, mean ± SEM. N indicates the number of determinations and n indicates the number of cells investigated from at least three different donors. See main text and/or the figure legends for further information. Statistical analyses were performed, using the Sigma Plot (v.13, Systat) software. Statistical significance was determined using the two-tailed t-test or the Holm Sidak one way ANOVA test, where indicated.

## Supporting information

all supplementary items

## Acknowledgments

We thank Dr. Vibor Laketa and the excellent service at the Infectious Disease Imaging Platform at the Center for Integrated Infectious Diseases at the Medical Faculty of Heidelberg University Hospital. We thank Sebastian Wurzbacher for performing scanning electron microscopy. We thank Marina Müller and Atdhe Kernaja for excellent technical assistance.

## Author contributions

Conceptualization: ML, USS, CPS, PP, MH, MK

Methodology and investigation: CPS, PP, S-YC, CK, LH, NK, MC, MK, ML

Data analysis: PP, CPS, NK, S-YC, CK, MK, ML

Supervision: ML, USS, MK, MH

Writing—original draft: ML

Writing—review & editing: all authors

## Data and materials availability

All data needed to evaluate the conclusions in the paper are present in the paper and/or the Supplementary Materials. Cryo-EM map of the spiral will be deposited to the electron microscopy data band and the accession code will be stated here.

