## Supplementary material for "KAHRP dynamically relocalizes to remodeled actin junctions and associates with knob spirals in *P. falciparum*-infected erythrocytes": all supplementary items

**This PDF file includes**

Tables S1 to S2

Figs. S1 to S13

Supplementary Materials and Methods

**Table S1. Antibodies and nanoprobes**

| Antibodies and nanoprobe | Vendor | dilutions |
|---|---|---|
| mAb Actin, clone AC40 (mouse) | Sigma-Aldrich | 1:100 |
| mAb Adducin α, clone 4D1 (mouse) | Santa Cruz Biotechnology | 1:100 |
| mAb Tropomodulin 1, OTI2C2 (mouse) | Novus Biologicals | 1:100 |
| mAb Spectrin ß, B-2 (mouse) | Santa Cruz Biotechnology | 1:100 |
| mAb Spectrin ß, A-12 (mouse) | Santa Cruz Biotechnology | 1:300 |
| Protein 4.1R (rabbit) | Sigma-Aldrich | 1:100 |
| Tropomyosin 5NM1, 5NM2 (sheep) | Merck | 1:100 |
| mAb Ankyrin-1, H-4 (mouse) | Santa Cruz Biotechnology | 1:200 |
| mAb KAHRP, 18.2 (mouse) | The European Malaria Reagent Repository | 0.8µg/ml |
| KAHRP$^{288-302}$ (rabbit) | Costum-made, Eurogentec | 1:500 |
| mAb 6xHIS, HIS-H8 (mouse) | Thermo Fisher | 1:100 |
| Star 580 goat anti-mouse | Abberior | 1:200 |
| Star 580 goat anti-guinea pig | Abberior | 1:200 |
| Star 580 goat anti-rabbit | Abberior | 1:200 |
| Star 580 donkey anti-sheep | Abberior | 1:200 |
| Star red goat anti-mouse | Abberior | 1:200 |
| Star red goat anti-guinea pig | Abberior | 1:200 |
| Star red goat anti-rabbit | Abberior | 1:200 |
| Alexa Fluor 647 goat anti-mouse | Thermo Fisher | 1:200 |
| $Ni^{2+}$-NTA-Atto 647 nanoprobe | Sigma-Aldrich | 1:400 |
| $Ni^{2+}$ NTA 5 nm gold | Nanoprobes | 1:30 |

NTA, Nitrilotriacetic acid

**Table S2. Oligonucleotides**

| **Primer name** | **sequence** |
|---|---|
| KAHRP-5-for | AGA ATA CTC GCG GCC ACT ATG AAA AGT TTT AAG AAC AAA AAT AC |
| KAHRP-3-rev | TTA CAA AAT GCT TAA CCG CGG TTA ACC ACA GCA TCC TCT TTT CTT C |
| KAHRP$^{rc}$-AflII-for | GGT CCA AAT ATA TTT GC*C TTA AG*G AAG AGA TTT CC |
| KAHRP$^{rc}$-AleI-rev | TTT TTT CTT TTC ACA GTC GTG GGA CTT ATG TTT TTT G |
| mEos-1x-AflII-for | GGT CCA AAT ATA TTT GCC TTA AGT GCG ATT AAG CCA GAC |
| mEos-1x-AflII-rev | TGG AAA TCT CTT CCT TCG TCT GGC ATT GTC AGG C |
| KAHRP-guide3-for | TAA GTA TAT AAT ATT TA TAG CGG CAG GTC CAC ATG GTT TTA GAG CTA GAA |
| KAHRP-guide3-rev | TTC TAG CTC TAA AAC CAT GTG GAC CTG CCG CTA TA AAT ATT ATA TAC TTA |
| KAHRP-guide4-for | TAA GTA TAT AAT ATT CAT GAA GGA AAT GAC GGT GA GTT TTA GAG CTA GAA |
| KAHRP-guide4-rev | TTC TAG CTC TAA AAC TCA CCG TCA TTT CCT TCA TG AAT ATT ATA TAC TTA |
| P1 | GTT TAT TTG AAC AAT ATT TAC TCC |
| P2 | ACC ACA GCA TCC TCT TTT CT |

Re-codonized KAHRP fragment between the AflII and AleI endo restriction sites

GCTTTGAGGAAGAGATTTCCATTGGGTATGAACGACGAAGATGAAGAAGGTAAAGAAGCTTTGGCCATCAAAGATAAGTTGCCAGGTGGTTTGGATGAATACCAAAATCAGTTGTACGGTATCTGTAACGAAACTTGTACTACTTGTGGTCCAGCTGCTATTGATTATGTTCCAGCTGATGCTCCAAATGGTTATGCTTATGGTGGTTCTGCTCATGATGGTTCACATGGTAATTTGAGAGGTCATGGTAACAAAGGTTCTGAAGGTTATGGTTATGAAGCTCCATACAATCCAGGTTTTAATGGTGCTCCAGGTTCAAATGGTATGCAAAATTACGTTCCACCACATGGTGCTGGTTATTCTGCTCCATATGGTGTTCCTCATGGTGCAGCTCATGGTTCTAGATATTCTTCATTCTCCTCCGTCAACAAATACGGTAAACATGGTGACGAAAAGCACCACTCTTCTAAGAAACATGAAGGTAATGATGGTGAGGGTGAAAAGAAGAAGAAGTCCAAAAAACACAAGGATCACGATGGCGAGAAAAAGAAGAGTAAAAAGCACAAAGATAACGAGGATGCCGAATCCGTTAAGAGCAAAAAACATAAGTCCCACGAC

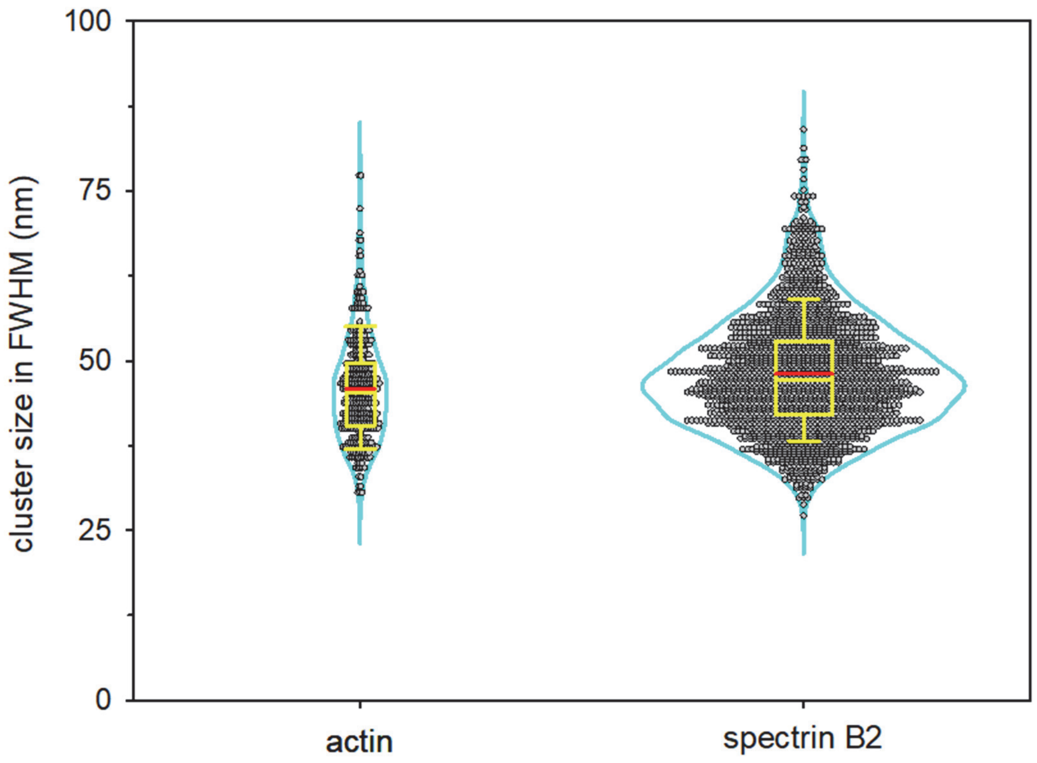


**Fig. S1. Cluster sizes in FWHM of actin and ß-spectrin N-terminus (spectrin B2) of uninfected erythrocytes.** Violin plots consisting of the Kernal density (cyan), the original data points and a box plot analysis. The box plots show the median (yellow horizontal line), the mean (red horizontal line) and the 25% and 75% quartile ranges. Error bars indicate the $10^{th}$ and $90^{th}$ percentile. Actin: N=357, n=23; N-terminus of ß-spectrin: N=2220, n=61.

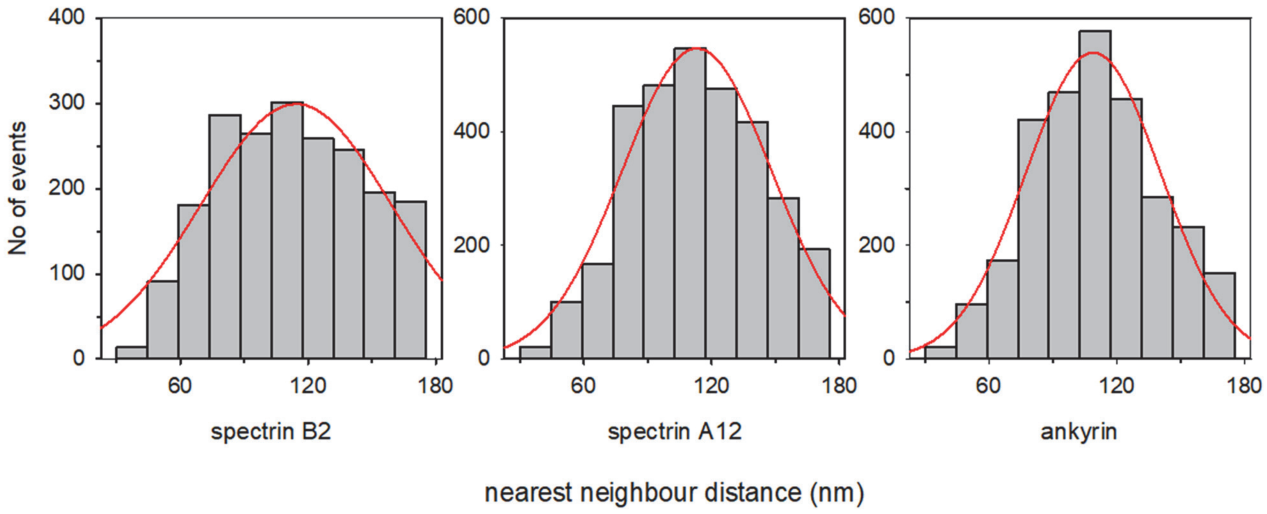


**Fig. S2. Histograms showing the nearest neighbor distance of ß-spectrin N-terminus (spectrin B2), ß-spectrin C-terminus (spectrin A12) and ankyrin.** A Gaussian function was fit to the data (red line).

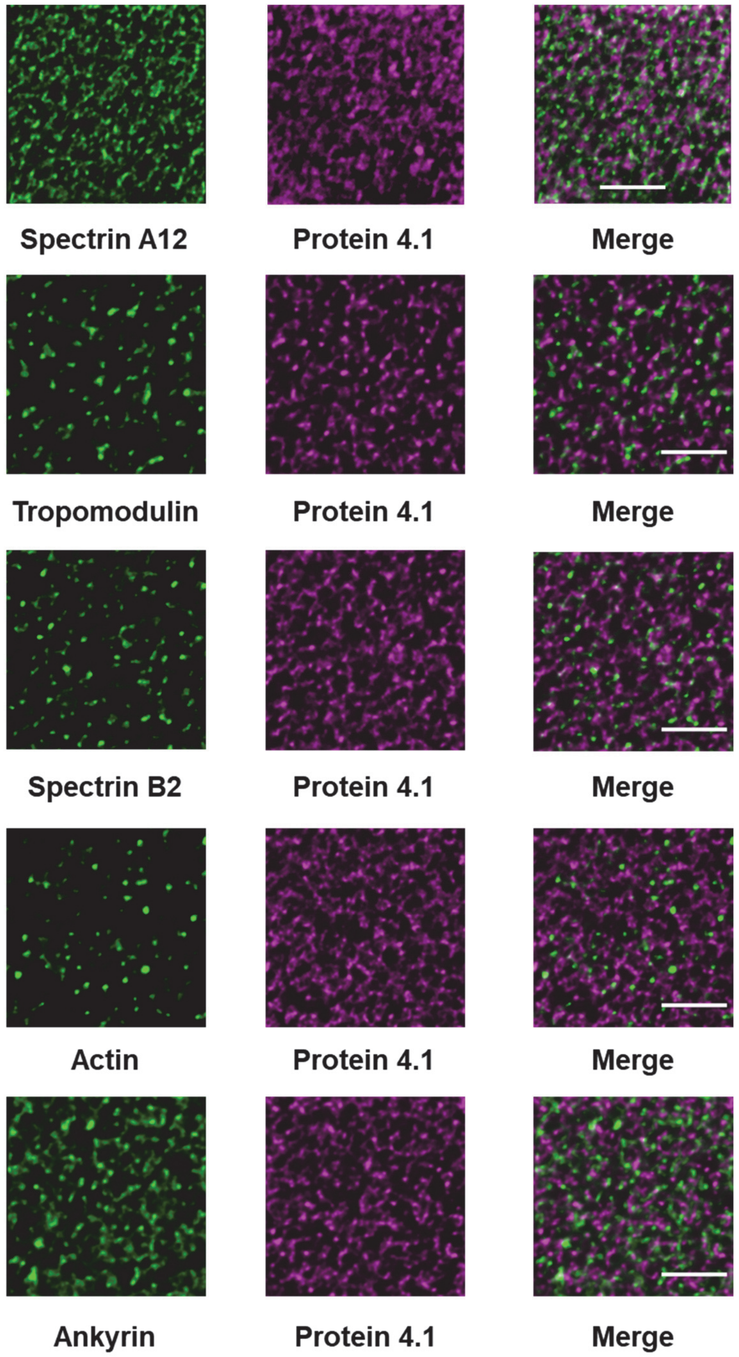


**Fig. S3. Representative STED images of exposed membrane skeletons from uninfected erythrocytes stained with antibodies to the targets indicated.** The two channels and the overlay are shown. Scale bar, 1 µm.

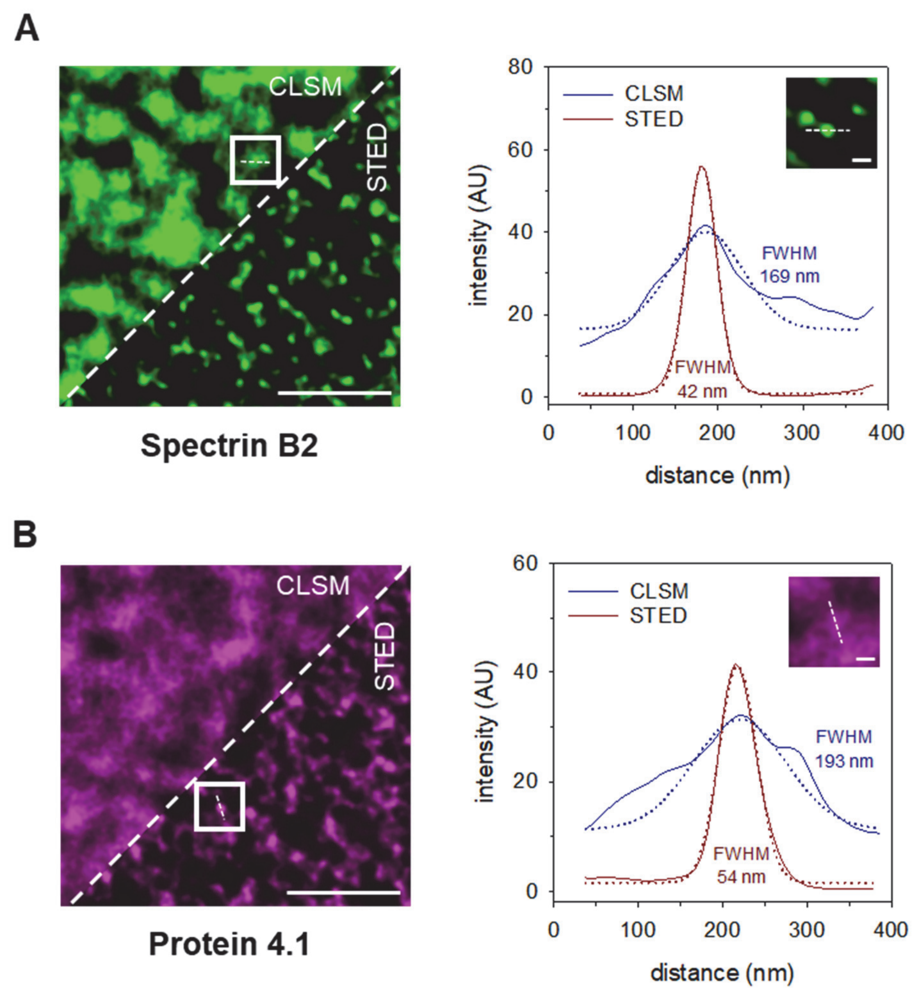


**Fig. S4. Comparison of confocal laser scanning microscopy (CLSM) and STED images of exposed membranes prepared from uninfected erythrocytes and stained with antibodies against the N-terminus of ß-spectrin (spectrin B2) and protein 4.1R.** (**A**) Split image showing CLSM and STED image. The following antisera were used: mouse monoclonal antibody spectrin B2 and goat anti-mouse antibody conjugated with Abberior star 580. Scale bar, 1 µm. White box and dashed line indicate the position of line profiles shown on the right. Gaussian fits to the intensity profiles (solid lines) are shown as dashed lines. Inset shows the STED image corresponding to the boxed area. Scale bar, 50 nm. (**B**) As in A., using a rabbit anti-protein 4.1R antiserum and an Abberior star red conjugated goat anti rabbit antiserum. The inset shows the CLSM image corresponding to the boxed area.

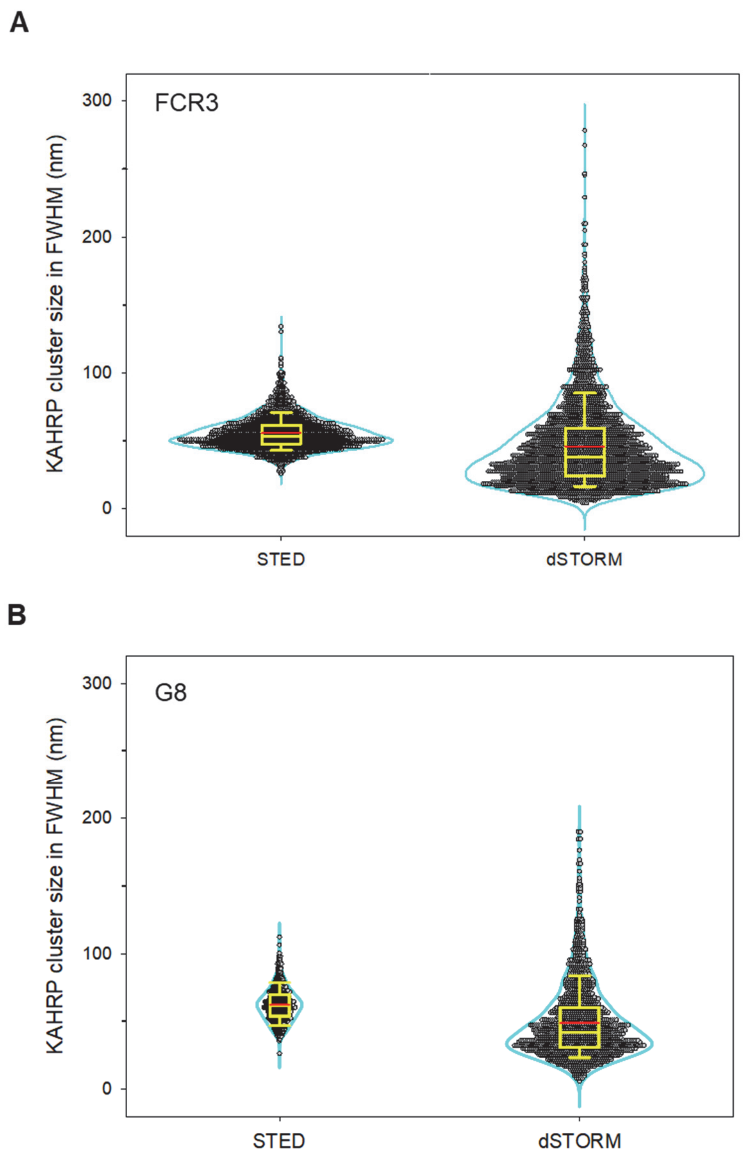


**Fig. S5. KAHRP cluster sizes in FWHM of the parental line FCR3 (A) and the KAHRP/mEOS-expressing mutant G8 (B), as determined by STED and dSTORM microscopy.** Violin plots consisting of the Kernal density (cyan), the original data points and a box plot analysis. The box plots show the median (yellow horizontal line), the mean (red horizontal line) and the 25% and 75% quartile ranges. Error bars indicate the $10^{th}$ and $90^{th}$ percentile. FCR3: N=2800, n=267 for STED and N=3383, n=14 for dSTORM. G8: N=646, n=29 for STED and N=1708, n=30 for dSTORM.

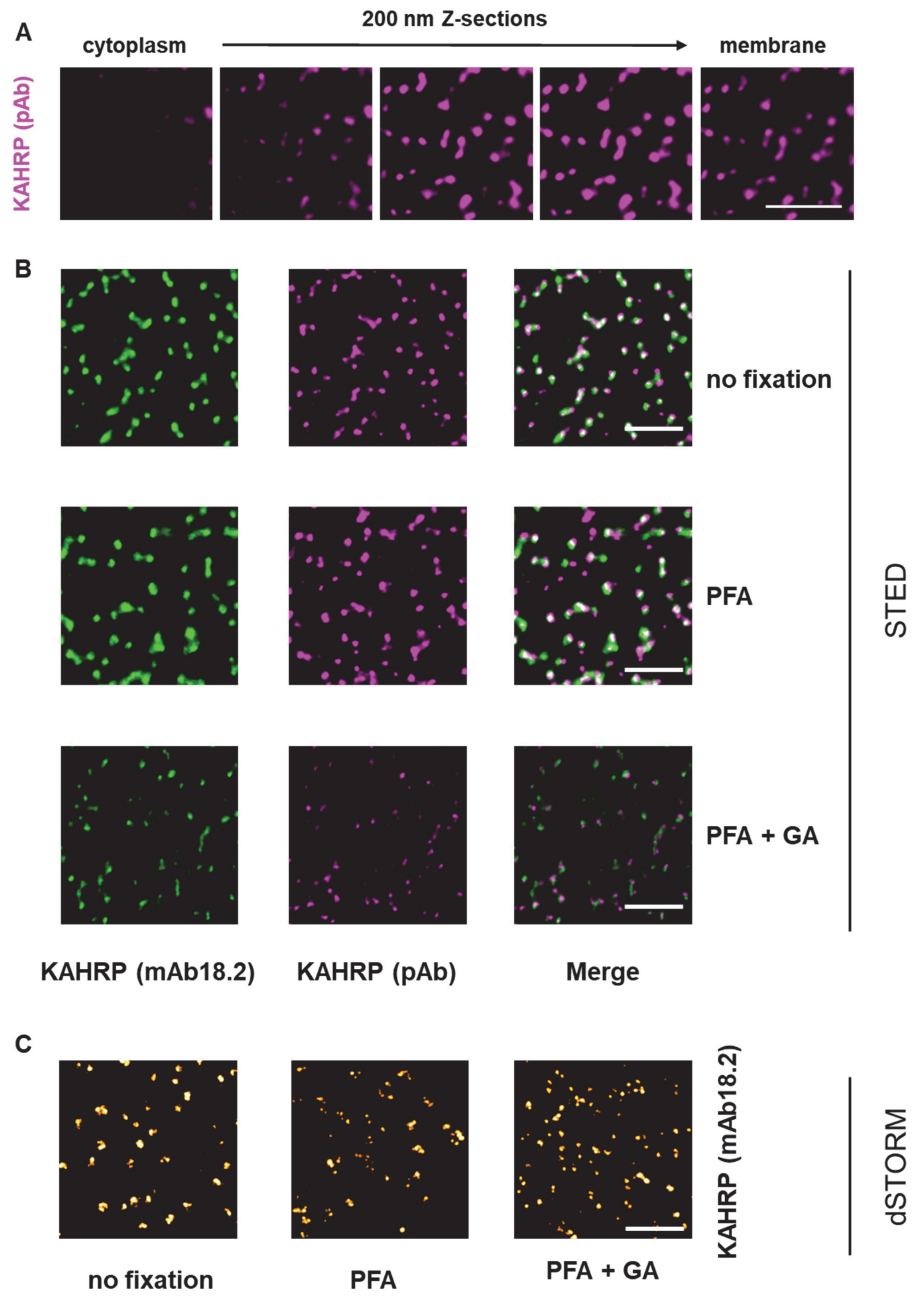


**Fig. S6. Super-resolution imaging of KAHRP under different fixation conditions.** (**A**) 3D STED imaging of exposed trophozoite membranes, using the anti-KAHRP peptide antiserum. 200 nm z-sections are shown. Scale bar, 0.5 µm. (**B**) Exposed trophozoites membranes were unfixed, fixed with 4% paraformaldehyde (PFA), or fixed with 4% paraformaldehyde and 0.0065% glutaraldehyde (PFA + GA) for 15 min before the sample was stained with the monoclonal anti KAHRP antibody mAB18.2 and the anti-KAHRP peptide antiserum. Representative STED images are shown. Scale bar, 1 mm. (**C**) Representative dSTORM images of unfixed and fixed anti-KAHRP stained (mAB18.2) exposed membranes from trophozoites. Scale bar, 1 µm.

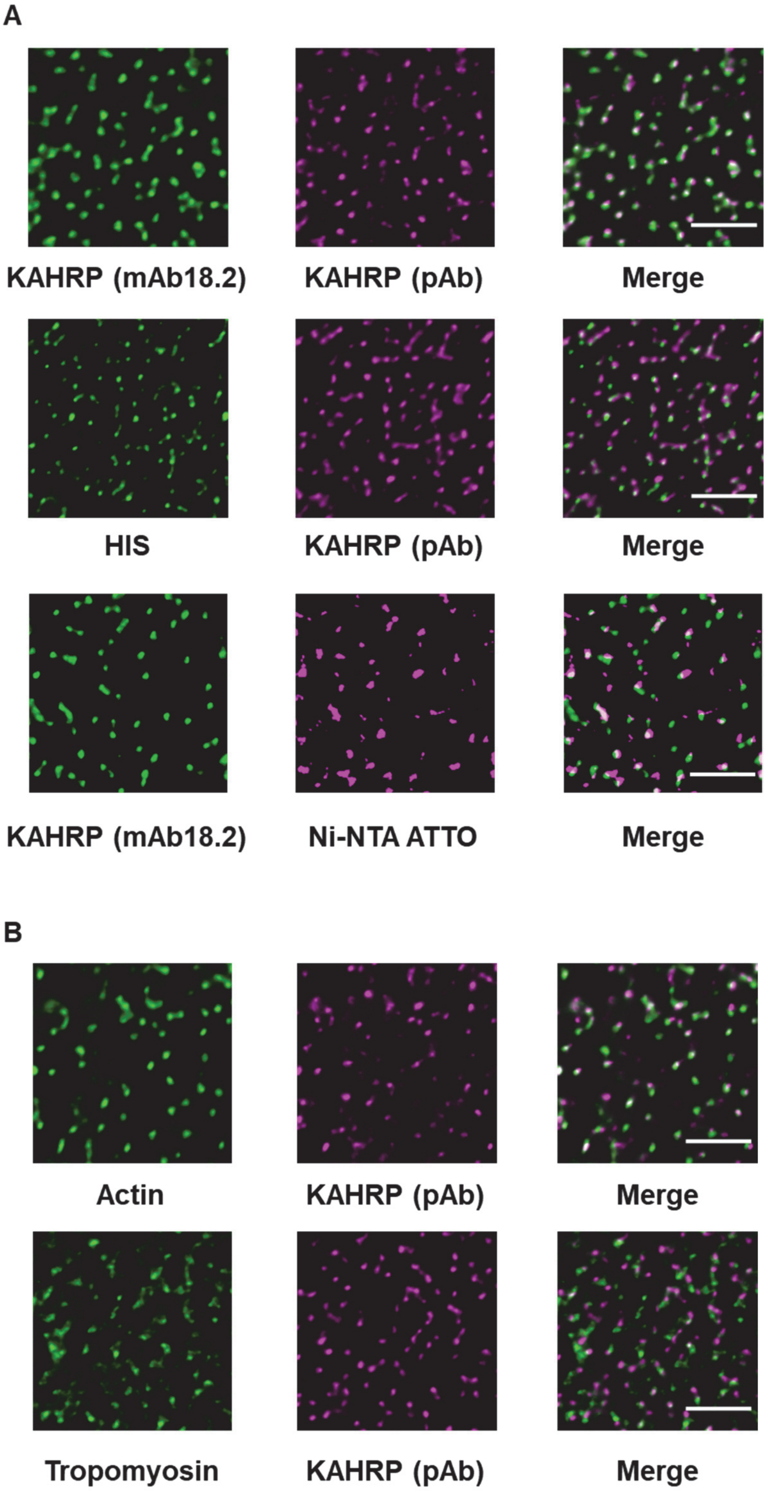
A
KAHRP (mAb18.2)
KAHRP (pAb)
Merge
HIS
KAHRP (pAb)
Merge
KAHRP (mAb18.2)
Ni-NTA ATTO
Merge
B
Actin
KAHRP (pAb)
Merge
Tropomyosin
KAHRP (pAb)
Merge

C

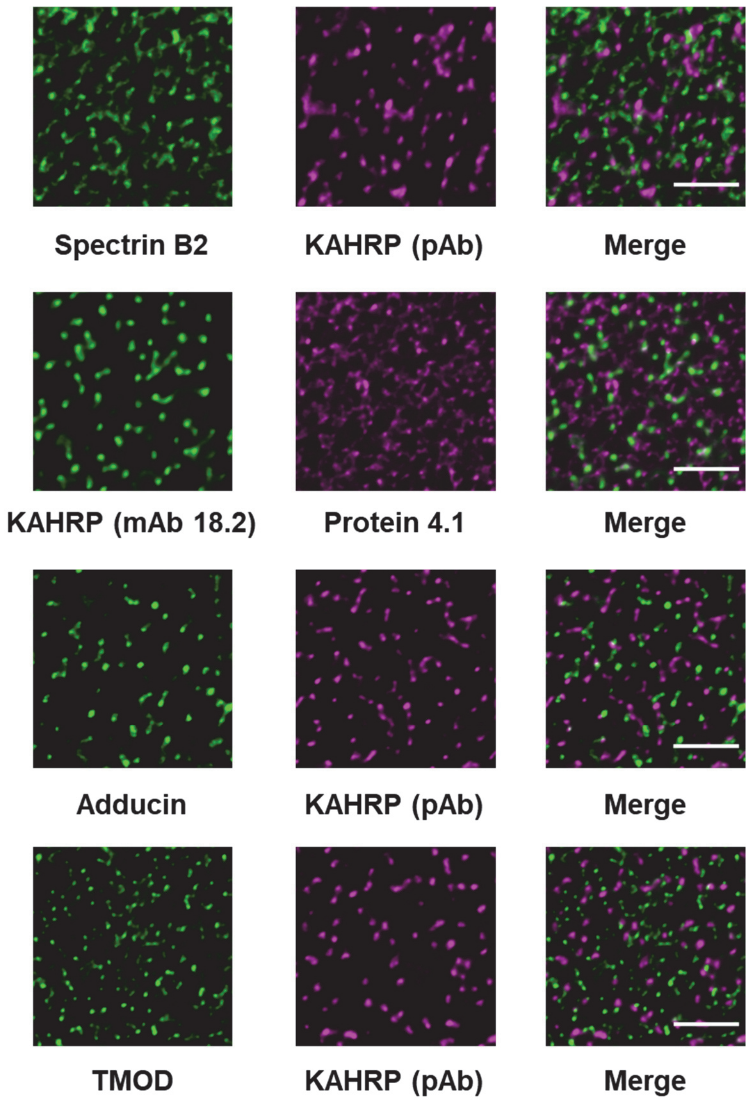
Spectrin B2
KAHRP (pAb)
Merge
KAHRP (mAb 18.2)
Protein 4.1
Merge
Adducin
KAHRP (pAb)
Merge
TMOD
KAHRP (pAb)
Merge

D

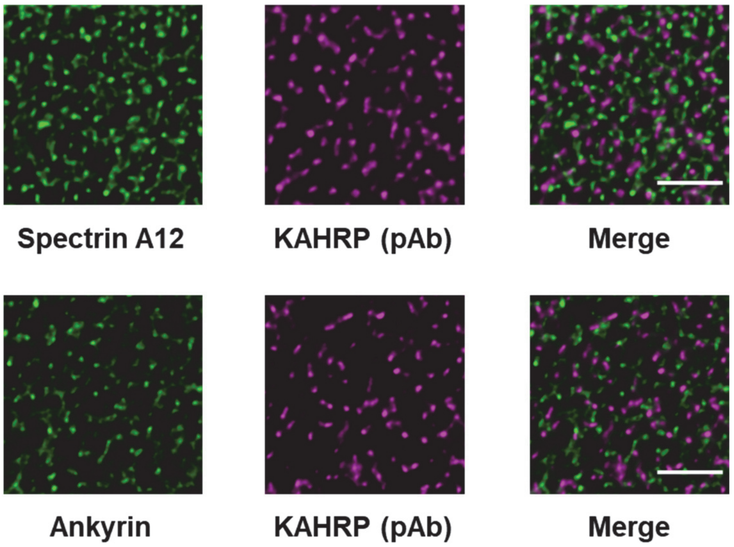
Spectrin A12
KAHRP (pAb)
Merge
Ankyrin
KAHRP (pAb)
Merge

**Fig. S7. Two-color STED images of exposed membranes prepared from trophozoites, showing KAHRP and a component of the membrane skeleton.** Separate STED images of the two channels and an overlaid STED image are shown. Scale bar, 1µm. (**A**) Various probes targeting KAHRP, including the monoclonal antibody mAB18.2, the peptide antiserum pAB, a monoclonal anti-histidine antibody (His), and a $Ni^{2+}$-NTA-ATTO nanoprobe. (**B**) Targeting KAHRP (using the peptide antiserum pAB) and actin and tropomyosin, two components of the actin junctional complex. (**C**) Targeting KAHRP (using the antiserum indicated) and components of the actin junctional complex, including the N-terminus of ß-spectrin (spectrin B2), protein 4.1R, adducing and tropomodulin (TMOD). (**D**) Targeting KAHRP and the C-terminus of ß-spectrin (spectrin A12) and ankyrin, two components of the ankyrin bridge.

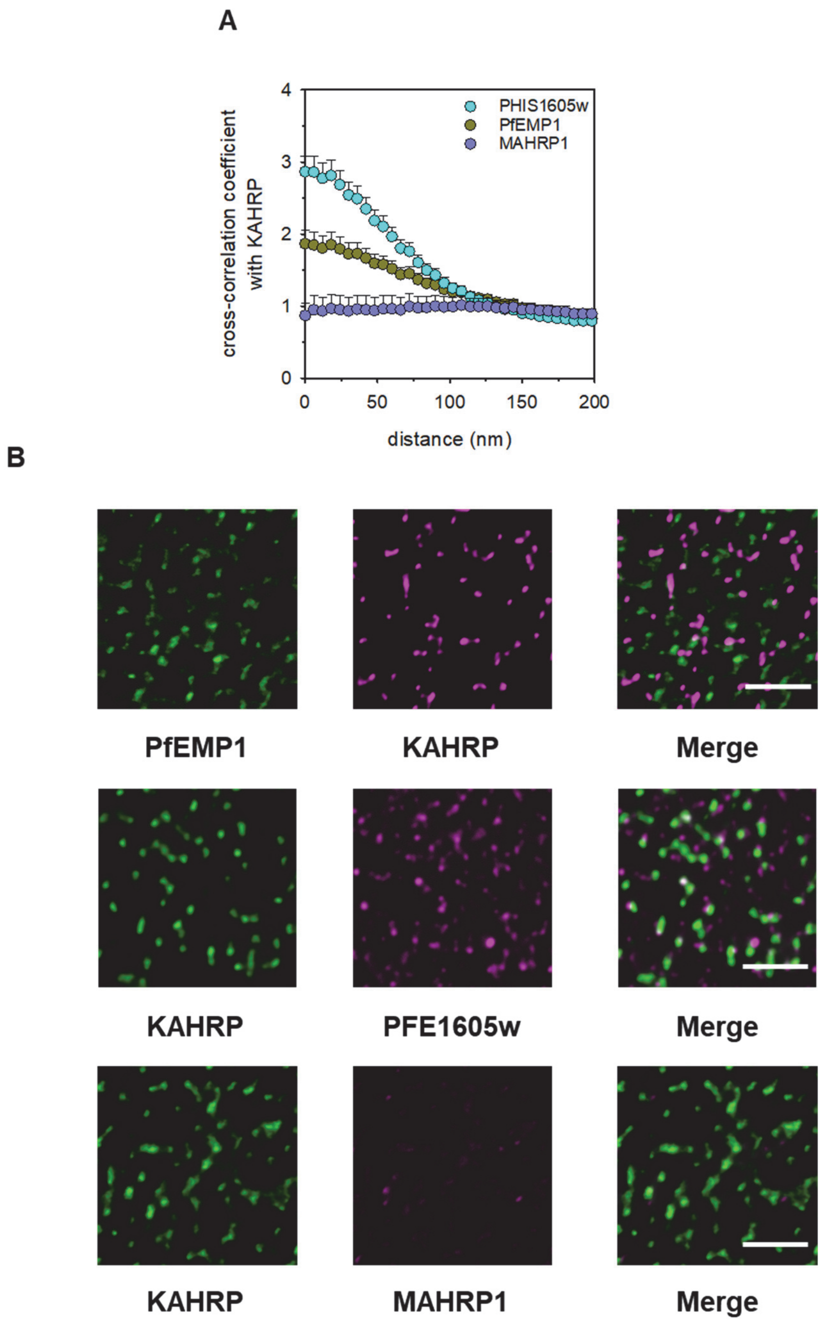


**Fig. S8. Co-localization of KAHRP with PfEMP1 and PHIS1605w and not MAHRP1.** (**A**) Calculated two-dimensional cross-correlations between KAHRP (using pAB) and PHIS1605w (cyan; n=26), PfEMP1 (olive green; n=15) and MAHRP1 (violet; n=14). The means ± SEM of n cells from at least three different donors are shown. (**B**) Two-color STED images of exposed membrane skeletons from trophozoite labeled with antisera against KAHRP, PfEMP1, PHIS1605w and MAHRP1. Separate STED images of the two channels and an overlaid STED image are shown. Scale bar, 1µm

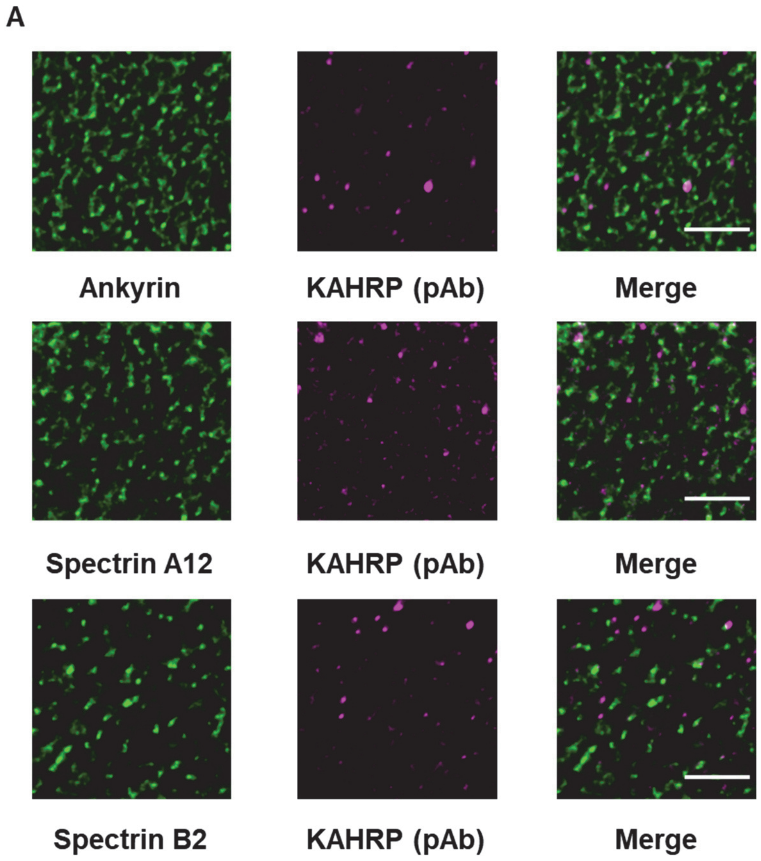
A
Ankyrin
KAHRP (pAb)
Merge
Spectrin A12
KAHRP (pAb)
Merge
Spectrin B2
KAHRP (pAb)
Merge

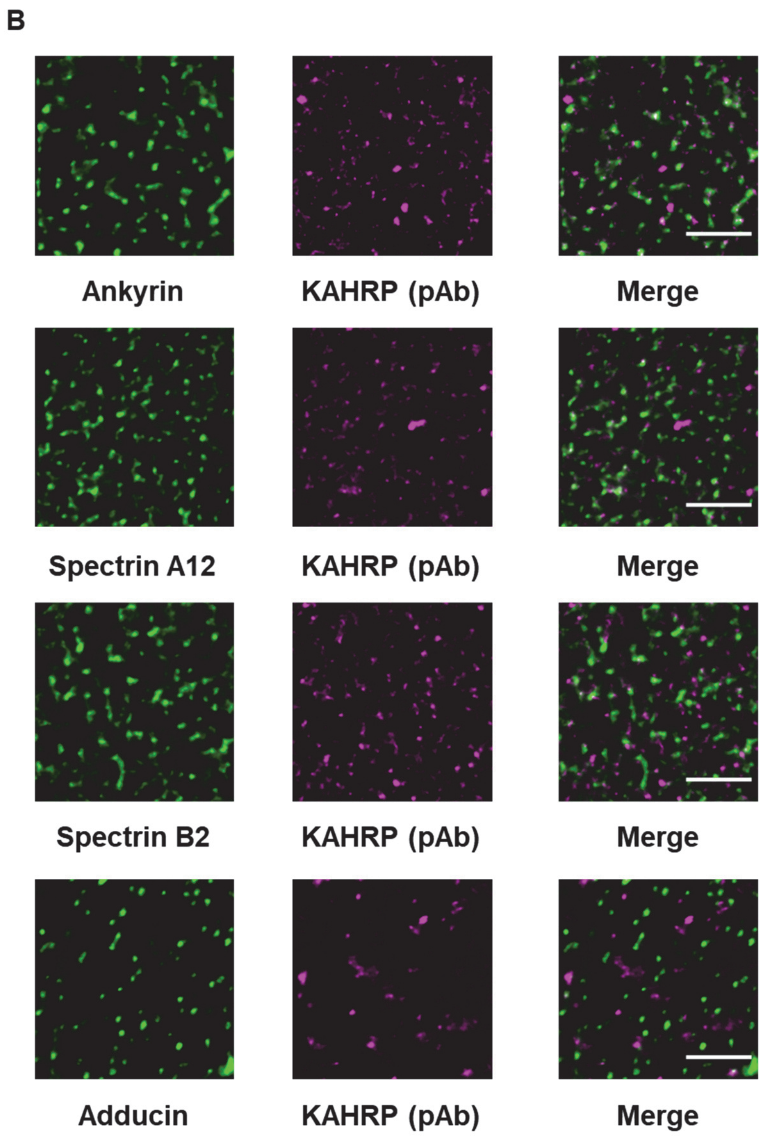
B
Ankyrin
KAHRP (pAb)
Merge
Spectrin A12
KAHRP (pAb)
Merge
Spectrin B2
KAHRP (pAb)
Merge
Adducin
KAHRP (pAb)
Merge

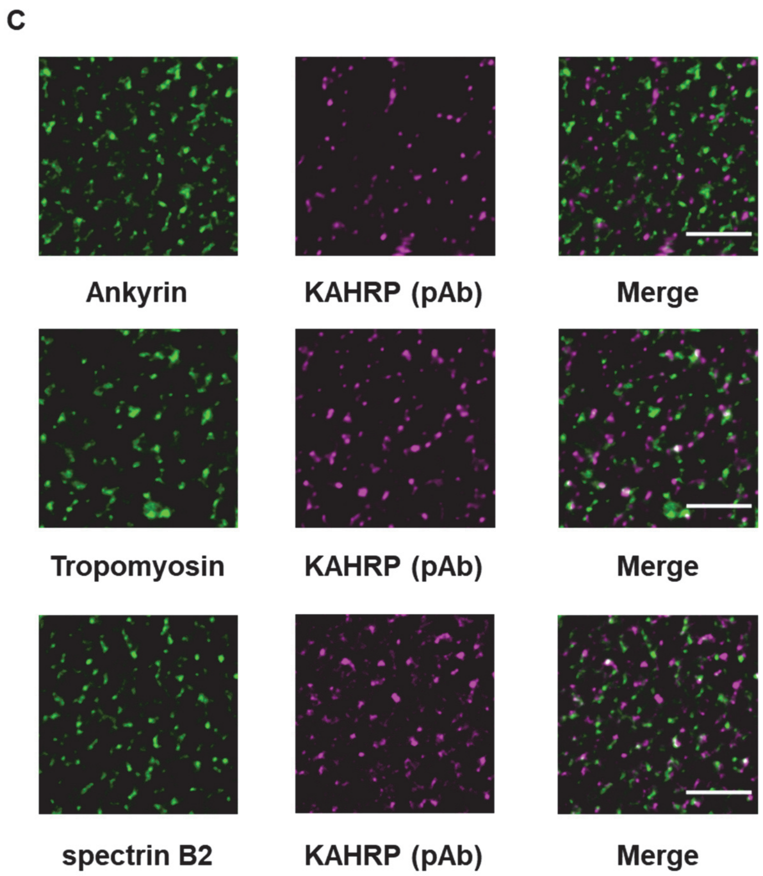


**Fig. S9. Two-color STED results of exposed membranes prepared from highly synchronized parasite cultures.** Separate STED images of the two channels and an overlaid STED image are shown. Scale bar, 1µm. (**A**) Exposed membranes were prepared from highly synchronized parasite cultures 12 ± 2 hours post invasion and stained with antisera against KAHRP, ankyrin, N-terminus ß-spectrin (spectrin B2) and C-terminus of ß-spectrin (spectrin A12). (**B**) as in A but investigating parasites 16 ± 2 hours post invasion. (**C**) as in A but investigating parasites 20 ± 2 hours post invasion

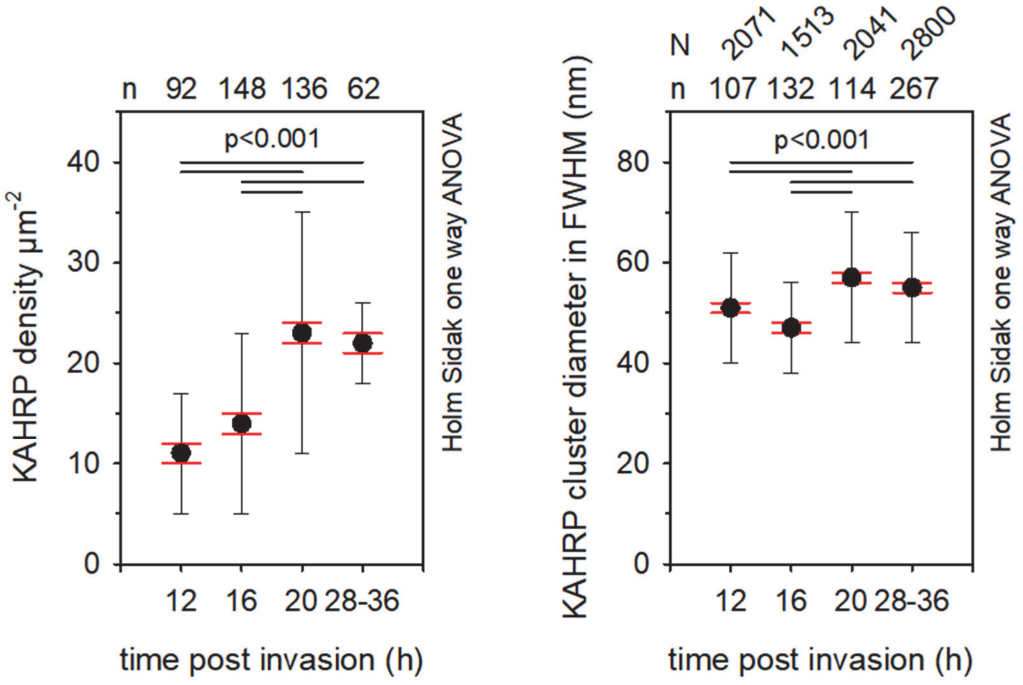


**Fig. S10. KAHRP signal density and size in FWHM as a function of time post invasion.** The means ± SEM (red whiskers) and SD (black whiskers) of N determinations from n cells are shown. Values for N and n are provided above the respective graphs. Statistical significance was assessed using the Holm Sidak one way ANOVA test.

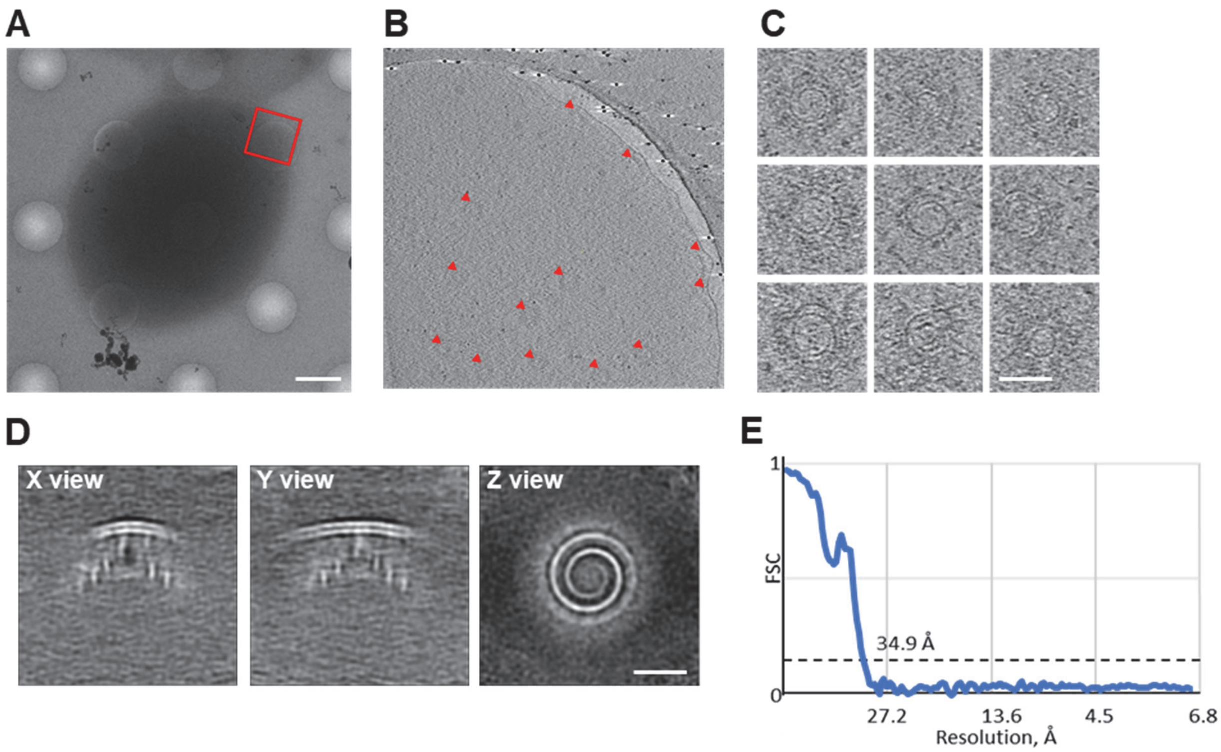


**Fig. S11. Cryo-tomography of knob spiral.** (**A**) Low magnification cryo-image of a membrane ghost prepared from a trophozoite infected erythrocyte. The image shows the ghost deposited on a holey EM-grid (Quantifoil 2/1). Scale bar: 2 µm. (**B**) An area for automated data acquisition is shown. Knobs and knob spirals in top views and side views are marked by red triangles. Scale bar, 200 nm. (**C**) Examples of top views of spirals. Scale bar, 50 nm. (**D**) Subtomogram averaging results with 35 Å bandpass filter. Scale bar, 20 nm. (**E**) Fourier Shell Correlation indicated a resolution of the reconstruction of 35 Å.

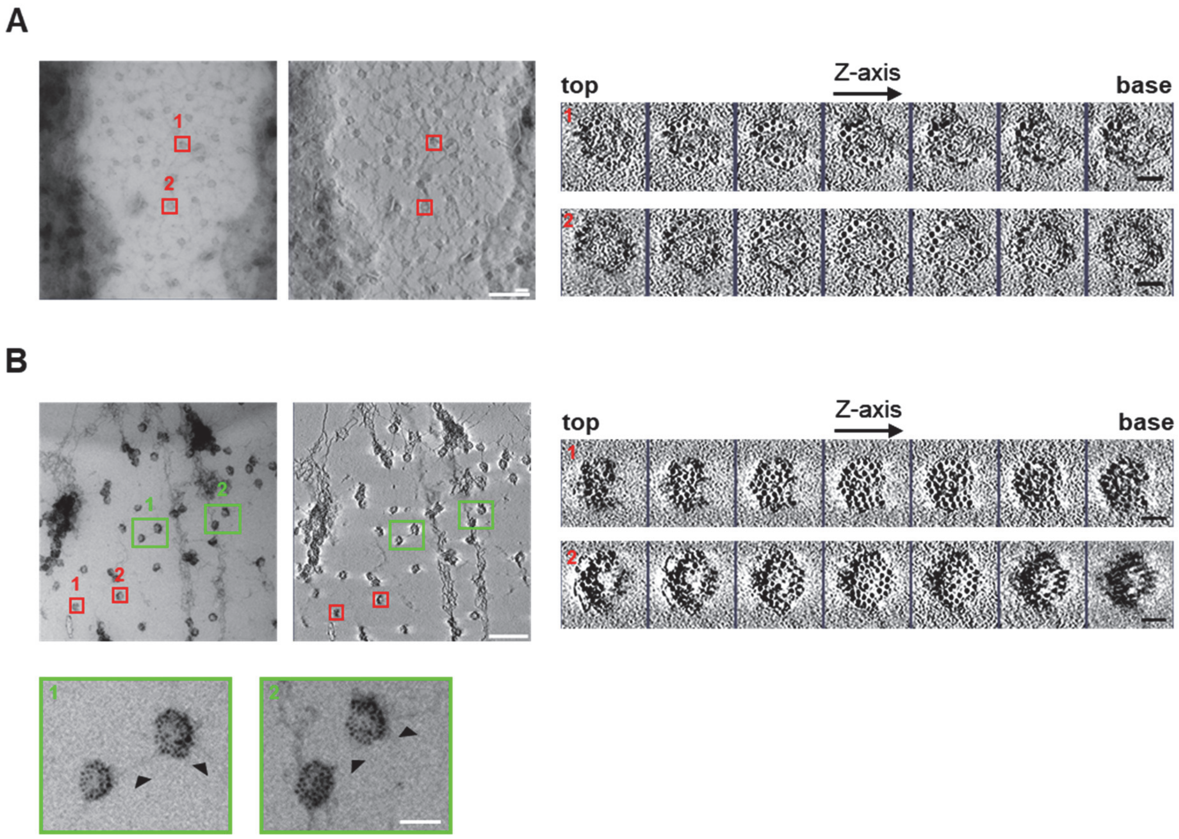


**Fig. S12. Immuno-EM labeling of the knob spiral with a $Ni^{2+}$-NTA 5 nm gold nanoprobe.** (**A**) Methyl cellulose embedded trophozoite membrane ghosts extracted with 0.2% NP40 Substitution and labelled with the $Ni^{2+}$-NTA 5 nm gold nanoprobe. Left: Transmission electron microscopic image (left image) and the corresponding tomographic reconstruction. Right: consecutive z-sections from an EM-tomogram showing labeling of the knob spiral scaffold with the $Ni^{2+}$-NTA 5 nm gold nanoprobe. The boxes indicate representative extracted volumes that were subsequently displayed in z-sections. The sections are shown from left to right from the top of the spiral to the base. Scale bars, 1 µm main images and 30 nm sub-images. (**B**) As in A, but analyzing trophozoite membrane ghost extracted with 1% NP40 Substitution. Green boxes: zoomed-in areas highlighting filaments extending from the spiral scaffold, as indicated by arrowheads. Scale bar, 50 nm

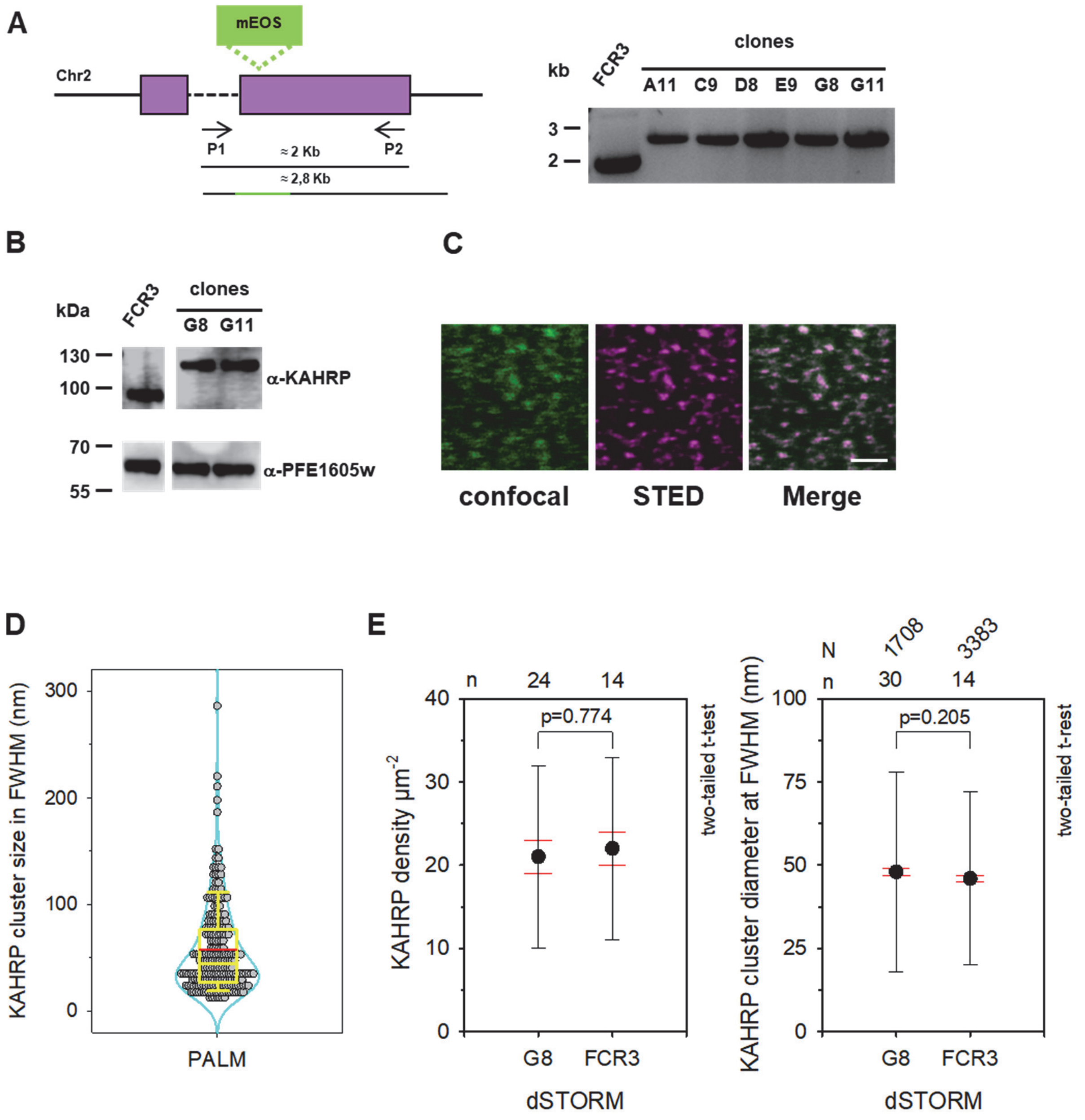


**Fig. S13. Generation and analysis of a KAHRP/mEOS2 expressing *P. falciparum* mutant line.** (A) Genomic organization of the *kahrp* locus on chromosome 2. The coding sequence of mEOS2 is indicated, as is the integration site. Relevant primers used for analysis of genetically altered lines and corresponding fragment lengths are shown. The phaerogram shows the result of a PCR analysis of genomic DNA prepared from the parental line FCR3 and five clonal mutant lines, using primers P1 and P2. The PCR products were sequenced and the insertion of the mEOS2 coding sequence in the *kahrp* coding sequence was confirmed in all mutant lines. A size marker is indicated in kb. (**B**) Western analysis using the anti-KAHRP peptide antibody (pAB). Note the shift in KAHRP signal in the investigated mutant lines G8 and G11, as compared with the parental line FCR3. An anti-serum to PFE1605w was used as a loading control. A size marker is indicated in kDa. (**C**) Imaging an exposed membrane prepared from the KAHRP/mEOS2 mutant G8 by correlative confocal fluorescence microscopy (left image; excitation wavelength, 488 nm and emission wavelength, 500-550 nm bandpass filter) and STED microscopy (STED) using the anti-KAHRP monoclonal mouse antibody mAB18.2 and an anti-mouse starred conjugated antibody (excitation wavelength, 640 nm and emission wavelength, 685-670 nm bandpass filter). The overlay of both images is shown. Scale bar, 1 µm. (**D**) KAHRP cluster size in FWHM of the KAHRP/mEOS-expressing mutant G8 (B), as determined by PALM. Kernal density pots with the original data points are shown. The box plots show the median (yellow horizontal line), the mean (red horizontal line) and the 25% and 75% quartile ranges. N=237, n=15. (**E**) Comparison of KAHRP cluster size in FWHM and distribution density from G8 and the parental line FCR3, as determined using dSTORM. The means ± SEM (red whiskers) and SD (black whiskers) of N determinations from n cells are shown. Values for N and n are provided above the respective graphs. Statistical significance was assessed using a two-tailed t-test.

Supplementary Materials for
KAHRP dynamically relocalizes to remodeled actin junctions and associates with knob spirals in P. falciparum-infected erythrocytes

Cecilia P. Sanchez, Pintu Patra, Shih-Ying Chang, Christos Karathanasis, Lukas Hanebutte, Nicole Kilian, Marek Cyrklaff, Mike Heilemann, Ulrich S. Schwarz, Mikhail Kudryashev, and Michael Lanzer*

##### Pair cross-correlation function

To compute cross-correlation between a pair of images, we follow the following step. First, we define two dimensional pair distance distribution (PDD) $P(r)$ for two images ($I_R(x,y)$, red channel and $I_G(x,y)$, green channel) [1, 4] as

$$P(r) = \frac{\sum_{i,j}\sum_{m,n} I_R(x_i,y_j)I_G(x_m,y_n)\delta(|\mathbf{r_{ij}} - \mathbf{r_{mn}}| - r)}{\sum_{i,j}\sum_{m,n} I_R(x_i,y_j)I_G(x_m,y_n)}. \tag{1}$$

A histogram of pair distance distribution is computed by using bins of width $\Delta r$ as

$$P(r, r+\Delta r) = \sum_{\rho=r}^{\rho=r+\Delta r} P(\rho). \tag{2}$$

Next, we radially average the resultant distribution to account for the increase in the area of radial bins. Specifically, we use fractional area, i.e., the area of each bin with respect to the image area, as the normalization factor N(r) [4, 6]. It is given by

$$N(r) = \frac{\pi\Delta r(2r+\Delta r)}{A_{image}}. \tag{3}$$

This normalization factor makes the distribution dimensionless and analogous to the definition of cross-correlation function for localization points data [3, 6]. The resultant distribution is the cross-correlation distribution (PCC) for two images,

$$C(r, r+\Delta r) = \frac{A_{image}}{\pi\Delta r(2r+\Delta r)} \sum_{\rho=r}^{\rho=r+\Delta r} \frac{\sum_{i,j}\sum_{m,n} I_R(x_i,y_j)I_G(x_m,y_n)\delta(|\mathbf{r_{ij}} - \mathbf{r_{mn}}| - \rho)}{\sum_{i,j}\sum_{m,n} I_R(x_i,y_j)I_G(x_m,y_n)}. \tag{4}$$

The above expression can be reduced for localization points data by using $I_R = \sum_{k=1}^{n_R}\delta(\mathbf{r_i} - \mathbf{r})$, the localisation points in red and $I_G = \sum_{k=1}^{n_G}\delta(\mathbf{r_j} - \mathbf{r})$, the localisation points in green. This leads to the expression PCC as

$$C(r, r+\Delta r) = \frac{\sum_{\rho=r}^{\rho=r+\Delta r}\sum_{i=1}^{n_R}\sum_{j=1}^{n_G}\delta(|\mathbf{r_i} - \mathbf{r_j}| - \rho)}{\pi\Delta r(2r+\Delta r)\eta}, \quad \eta = \frac{n_R \times n_G}{A_{image}} \tag{5}$$

The numerator is evaluated by making a histogram of pairwise distances of single-molecule localization points from two images. The denominator is calculated by repeating the same procedure with an equal number of randomly distributed points of two colors in the same area. The average distance distribution for randomly distributed data-set is given by $\pi\Delta r(2r+\Delta r)\eta$, where $\eta$ is the density of pair-wise distances [4, 6]. This method has been used in the recent works [3–6] for super-resolution microscopy localization data-sets.

##### Improving pair distance distribution using image re-sampling

The first step for estimation cross-correlation is the numerical calculation of pair distance distribution (PDD) from images. We notice PDD at short distances is limited by the pixel dimension of the image, e.g., the minimum separation distance can be 1 pixel. To estimate PDD at a smaller distance, we interpolate the images to a finer scale using image interpolation or re-sampling. This procedure does not lose any information

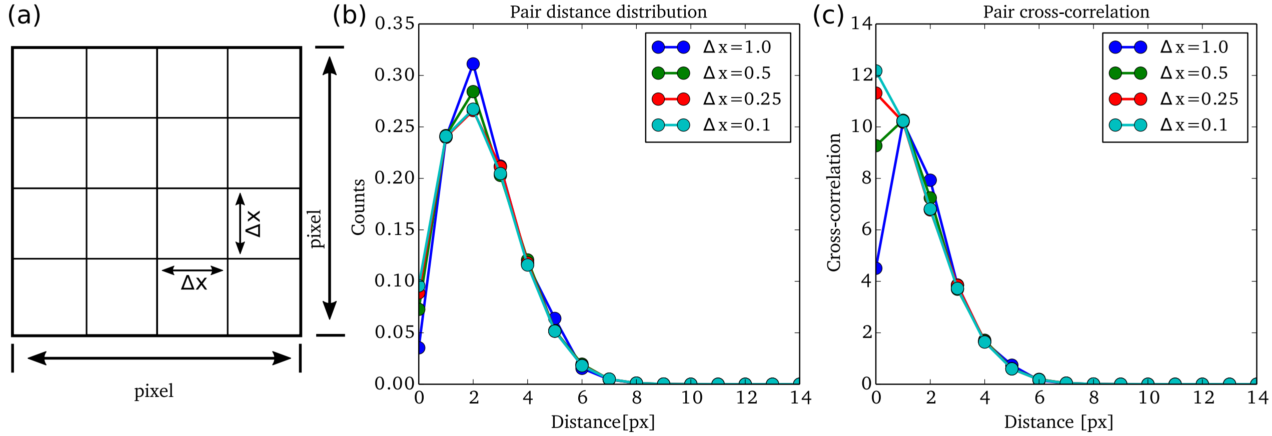


FIG. 1: (a) Each image pixel in the original image is subdivided into multiple smaller pixels with an area $\Delta x \Delta x$. (b) Normalised pair-distance distribution for two 2D-guassian separated by a distance of 0 pixel. (c) Pair cross-correlation function for two 2D-guassian separated by a distance 0 px for different $\Delta x$ values.

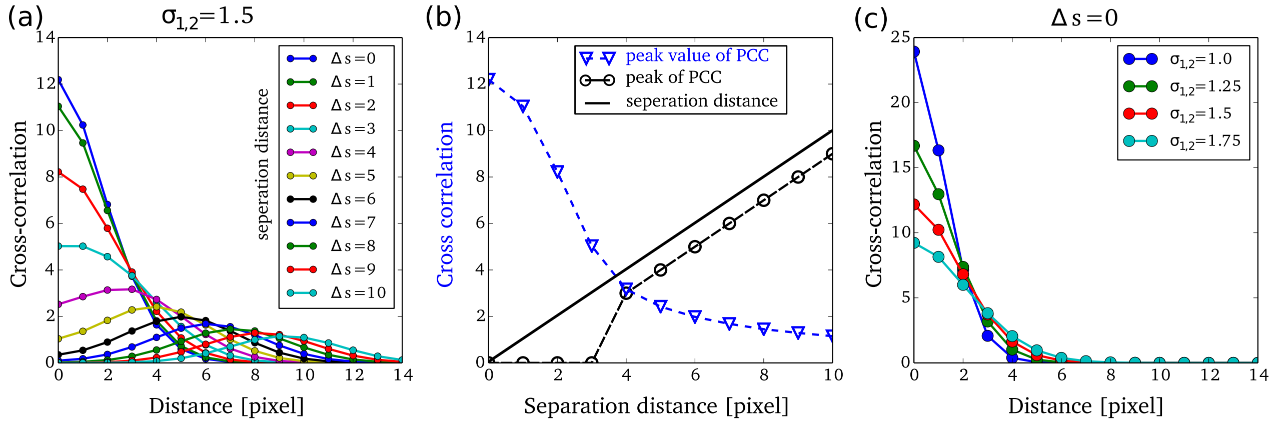


FIG. 2: (a)Pair cross-correlation distribution for different separation distances ($\Delta s$) of the signal pairs. (b) Separation estimated from the peak distance value in PCC and corresponding PCC value. (c) Pair cross-correlation distribution for different standard deviation of the two signals for zero separation.

as a single fluorescence signal covers multiple pixels. Specifically, we divide each pixel area into smaller grids (Fig. 1a) and interpolate the signal intensity using bi-linear interpolation. The improvement from this step saturates as the grid size becomes smaller, as shown in Fig. 1b. We choose $\Delta x = 0.1$ pixel for all the experimental images. The cross-correlation distribution also improves due to the improvement in PDD. The peak in cross-correlation shifts to zero from a one-pixel distance (Fig 1c).

The pair cross-correlation function is highest at zero inter-molecular distance for smaller separation (particularly when the separation distance $d$ is comparable to $\sqrt{\sigma_1^2 + \sigma_2^2}$, where $\sigma_{1,2}$ are the standard deviations for the two signals [1, 2]). The cross-correlation value at zero distance increases as the inter-molecular distance decreases (Fig. 2a). At the large separation distances, the pair correlation function shows a peak at finite distance (Fig. 2b).

Another factor that influences the cross-correlation value is the noise in fluorescence signals. Increase in the standard deviation in signals decreases cross-correlation value (Fig. 2c). Physically, this means sharper signals will lead to higher cross-correlation values compared to broader signals for a fixed separation distance. For our analysis, we choose a bindwidth $\Delta r$=0.4 pixel= 6 nm (1 pixel = 15 nm). The code for cross-correlation analysis is written in Python.

### Python code for computation of pair cross-correlation

```
import numpy as np
import matplotlib.pyplot as plt
from scipy import interpolate
from scipy.spatial.distance import cdist
from PIL import Image

ROIval=np.loadtxt('ROI.txt')#Input file for region of interest
Xmin = int(ROIval[0])
Ymin = int(ROIval[1])
Xmax = int(ROIval[2])
Ymax = int(ROIval[3])

filename1='STARRED'
filename2='594'
threshold=20 #Intensity below this value is considered as noise

img1 = Image.open(str(filename1)+'.tif')  #Loading image 1
image1 = np.array(img1) #Converting image in array
image1 = image1[0:,0:,0] #Selecting the right channel (RGB) of the image array: 0 for red
image1[image1<=threshold]=0 #Image thresholding
image1=image1[Ymin:Ymax,Xmin:Xmax] #Interested region is cropped out

#Same steps are repeated for Image 2
img2 = Image.open(str(filename2)+'.tif')
image2 = np.array(img2)
image2 = image2[0:,0:,1] #Selecting the right channel (RGB) of the image array: 1 for green
image2[image2<=threshold]=0
image2=image2[Ymin:Ymax,Xmin:Xmax]

#Image resampling or interpolation
x = np.arange(0, Xmax-Xmin, 1) #Existing x-grid
y = np.arange(0, Ymax-Ymin, 1) #Existing y-grid
f1 = interpolate.interp2d(x, y, image1, kind='linear')
f2 = interpolate.interp2d(x, y, image2, kind='linear')

dxy=0.1 #New grid spacing

xnew = np.arange(0, Xmax-Xmin, dxy) # New x-grid
ynew = np.arange(0, Ymax-Ymin, dxy) # New y-grid
image1 = f1(xnew, ynew) #Resampled image 1
image2 = f2(xnew, ynew) #Resampled image 2

image1new=image1.copy()
image2new=image2.copy()

loc1=np.array(np.where(image1new!=0)).T #Contains coordinates of the non-zero grids
val1=image1[np.where(image1new!=0)]*1.0 #Contains intensity values of such grids.
loc2=np.array(np.where(image2new!=0)).T
val2=image2[np.where(image2new!=0)]*1.0

dist12=cdist(loc1, loc2) #distance between all the locations
val=np.dot(np.array([val1]).T,np.array([val2])) #values of intenstiy product in these locations
distances=np.ravel(dist12*dxy) #putting all distances in a linear array
values=np.ravel(val) #putting the corresponding intensity produc in a linear array

lengthscale=15
binwidth=0.4
rMax=14

Sx=Xmax-Xmin #Width of the image
Sy=Ymax-Ymin #Height of the image
XY=(Sx*Sy) #Area of the image
Num=np.sum(val1)*np.sum(val2)# Values of all intensty product
CC=np.array([0,0])
for dd in range(0,int(rMax/binwidth)): #preparing histogram
    pair hist=np.sum(values[np.where((distances>=dd*binwidth) & (distances<dd*binwidth+binwidth))])/Num
    rand hist=3.14*binwidth*(2*dd*binwidth+binwidth)/XY
    CC=np.vstack([CC,[(dd+0.5)*binwidth*lengthscale,pair_hist/rand_hist]])

np.savetxt('PCC.dat', PCC,fmt='%2.2f  %2.4f')
```


[1] L Stirling Churchman, Zeynep Ökten, Ronald S Rock, John F Dawson, and James A Spudich. Single molecule high-resolution colocalization of cy3 and cy5 attached to macromolecules measures intramolecular distances through time. *Proceedings of the National Academy of Sciences*, 102(5):1419–1423, 2005.

[2] Stefan Niekamp, Jongmin Sung, Walter Huynh, Gira Bhabha, Ronald D Vale, and Nico Stuurman. Nanometer-accuracy distance measurements between fluorophores at the single-molecule level. *Proceedings of the National*

*Academy of Sciences*, 116(10):4275–4284, 2019.

[3] Leiting Pan, Rui Yan, Wan Li, and Ke Xu. Super-resolution microscopy reveals the native ultrastructure of the erythrocyte cytoskeleton. *Cell reports*, 22(5):1151–1158, 2018.

[4] Joerg Schnitzbauer, Yina Wang, Shijie Zhao, Matthew Bakalar, Tulip Nuwal, Baohui Chen, and Bo Huang. Correlation analysis framework for localization-based superresolution microscopy. *Proceedings of the National Academy of Sciences*, 115(13):3219–3224, 2018.

[5] Prabuddha Sengupta, Tijana Jovanovic-Talisman, Dunja Skoko, Malte Renz, Sarah L Veatch, and Jennifer Lippincott-Schwartz. Probing protein heterogeneity in the plasma membrane using palm and pair correlation analysis. *Nature methods*, 8(11):969, 2011.

[6] Matthew B Stone and Sarah L Veatch. Steady-state cross-correlations for live two-colour super-resolution localization data sets. *Nature communications*, 6:7347, 2015.
